# Bridging the gap – Spontaneous fluctuations shape stimulus-evoked spectral power

**DOI:** 10.1101/2020.06.23.166058

**Authors:** Soren Wainio-Theberge, Annemarie Wolff, Georg Northoff

## Abstract

Spontaneous fluctuations of neural activity have been shown to influence trial-by-trial variation in perceptual, cognitive, and behavioural outcomes. This implies that these fluctuations affect stimulus-related neural processes, and hence should affect stimulus-evoked neural activity. However, the mechanisms by which spontaneous neural activity shapes stimulus-evoked neural activity have rarely been examined. Employing a large-scale magnetoencephalographic dataset, as well as an electroencephalographic replication dataset, we observed that for high-frequency power, high pre-stimulus activity leads to greater evoked desynchronization (negative interaction); in contrast, for low-frequency power, high pre-stimulus activity induces greater event-related synchronization (positive interaction). We show that both positive and negative interactions are manifest primarily in cortical oscillations, rather than scale-free activity, and can also be observed in the time domain. In summary, we demonstrate positive and negative spontaneous-evoked interaction in multiple electrophysiological processes; these mechanisms “bridge the gap” between spontaneous and evoked activity and provide novel insights into how spontaneous activity influences behaviour and cognition.

## Introduction

Even in the absence of specific experimental stimulation, neural activity displays spontaneous fluctuations with a characteristic temporal and spatial structure^1–5^. This spontaneous neural activity has been associated with various forms of internally-oriented cognition, such as mind wandering^6–9^, self-referential processing^10–15^, mental time travel^16–19^ and social cognition/theory of mind^20,21^. While spontaneous activity is typically studied in stimulus-free “resting-state” designs, it persists in cognitive tasks as trial-by-trial fluctuations in neural activity. The main goal of our study is to “bridge the gap” between spontaneous and stimulus-evoked activity, by showing the neural mechanisms of their interaction.

The use of behavioural tasks to probe neural activity relies on the central assumption that changes in neural activity following the presentation of a stimulus are reflective of the brain’s processing of that stimulus. However, multiple recent studies have now demonstrated that spontaneous neural activity *prior* to stimulus onset can predict or influence the subject’s stimulus-related perception^22–29^, sense of self^30,31^, consciousness^28,32–39^, attention^40^, reaction time^41^, and working memory^42^. The pervasive influence of pre-stimulus activity on behavioural outcomes raises questions about the neural mechanism of this effect. If spontaneous pre-stimulus neural activity is to have behaviourally observable effects, then it must influence the brain’s processing of the stimulus, and hence also influence neural activity changes following stimulus onset. However, the mechanisms by which pre-stimulus spontaneous activity affects post-stimulus evoked activity remain unclear. In this paper, we will term this relationship *spontaneous-evoked interaction*.

While many different forms of spontaneous-evoked interaction are possible, one of the simplest to consider is a direct correlation between spontaneous activity in the prestimulus period and the magnitude of the stimulus-evoked response. In a *negative interaction scheme*^43^, spontaneous and evoked activity are negatively correlated, such that high spontaneous activity in the prestimulus period leads to a reduced stimulus-evoked increase in activity (or a greater stimulus-evoked decrease in activity). In contrast, a *positive interaction scheme* describes a positive correlation, where high prestimulus activity leads to greater stimulus-evoked increases. These models provide a useful framework to study the features and mechanisms of the interaction between spontaneous and task-evoked activity. Note, however, that correlation as a method is insufficient to detect these interactions, as activity in the poststimulus period reflects a mixture of evoked and spontaneous activity. Instead, more comprehensive methods have been developed for this purpose, which are described in the Results and Methods sections.

Previous small-sample fMRI studies have shown evidence for a negative interaction between spontaneous and evoked hemodynamic amplitudes^43,44^. However, given that fMRI reflects as-yet-unclear neuro-vascular coupling^45,46^, an investigation of the interaction between spontaneous and evoked activity in electrophysiology would be valuable in unraveling the neural processes involved. Electroencephalography (EEG) and magnetoencephalography (MEG) provide a window into an abundance of neurophysiological phenomena^47^, and analyses in the time and frequency domains reflect distinct neurophysiological processes. The time-domain electrophysiological signal is known to reflect synchronous postsynaptic potentials of many neurons^48^. In contrast, frequency domain analyses allow one to record cortical oscillations such as alpha (8-13 Hz), theta (4-8 Hz), and beta (13-25 Hz)^47^; these are thought to reflect cortical feedback loops or neurotransmitter-related processes^49,50^. Frequency-domain analyses also reveal arrhythmic “scale-free” activity, which has been associated with excitation-inhibition balance^51–53^ and complex network models of self-organized criticality^54–56^. It remains unknown which of these electrophysiological parameters, if any, shows an interaction between spontaneous and evoked activity.

In the present study, we investigated the interaction between spontaneous and stimulus-evoked neural activity across a diverse set of electrophysiological processes. For this purpose, we employed a large-scale MEG data set with a simple sensory paradigm^57,58^, as well as a replication EEG dataset with a more complex cognitive task^59^. In the frequency domain, we observed widespread interaction between spontaneous and evoked spectral power, a finding which was consistent across modalities and tasks. The type and magnitude of this interaction varied between frequency bands and was found to be most prominent in the dynamics of cortical oscillations, rather than scale-free activity. Interaction between spontaneous and evoked activity was also observed in the time domain, where these interactions appeared to be task- or modality-specific. Our study sheds new light on the interaction between spontaneous and evoked neural activity and the neurophysiological processes involved; in turn, this promises to explain the mechanism by which trial-by-trial fluctuations in spontaneous activity affect cognitive and perceptual outcomes.

## Results

Going beyond simple correlation and avoiding the ubiquitous phenomenon of regression to the mean^60^, the relationship between spontaneous activity and evoked activity can be investigated in two different ways, which we term “TTV method” and “pseudotrial method”. For the TTV method, trial-to-trial variability^43^ (TTV) is computed as the standard deviation of the signal across trials: according to the law of total variance (see Methods), a poststimulus reduction in TTV must be indicative of a negative interaction. Alternatively, one can calculate the influence of prestimulus activity in a more direct way using the pseudotrial method. Trials are split into above-median and below-median pre-stimulus activity level, and separate post-stimulus activity time courses are computed relative to these baselines. These estimates are then corrected by subtracting “virtual” trials, i.e., pseudotrials^43^ drawn from the intertrial interval (preceding the prestimulus period). This corrects for ongoing spontaneous fluctuations with the same initial conditions, and thereby corrects for regression to the mean of spontaneous activity.

If the trials with high prestimulus activity show a greater evoked increase (or smaller evoked decrease) than trials with low prestimulus activity, this is evidence for a positive interaction. If, by contrast, trials with high prestimulus activity show a smaller evoked increase (or greater evoked decrease) than trials with low prestimulus activity, then this is evidence for a negative interaction. As the pseudotrial method is capable of detecting both positive and negative interaction (and because of our simulation results – see supplementary figures S2 and S3), we focused our analysis on this method, using the TTV method to confirm these findings. Both methods were applied to time-domain and frequency-domain analyses. These methodologies and their theoretical justifications are described in detail in the methods.

### Spontaneous-evoked interaction in the frequency domain – effects vary by frequency band

The first aim of our study was to investigate spontaneous-evoked interaction in both the time and frequency domains. Following the observations in fMRI^43,44^, we hypothesized negative interaction between pre- and post-stimulus activity: that is, high pre-stimulus activity should lead to a stronger decrease (or weaker increase) in post-stimulus activity than low pre-stimulus activity. Given that fMRI signals are driven strongly by spectral power^61–65^, we hypothesized pre-post-stimulus correlation to be visible primarily in frequency-domain representations. Our second aim was then to assess the possible differences between frequency bands in their relationship between spontaneous and evoked activity, following previous findings of differential trial-to-trial variability reduction in different frequency bands^59,66^.

Using the second method, i.e., the method of pseudotrials, we first tested for the presence of interaction between spontaneous and evoked activity in MEG by comparing the effect of low vs. high pre-stimulus spectral power on post-stimulus event-related spectral perturbation (ERSP; Figure 2). In the broadband data, we observed a significant interaction beginning at around 200 ms poststimulus (p = 0.002). With respect to specific frequency bands, our results show significant spontaneous-evoked interactions in multiple bands including delta (p = 0.002), theta (p = 0.002), alpha (p = 0.002), beta (p = 0.002), and low gamma bands (p = 0.002).

**Figure 1.**
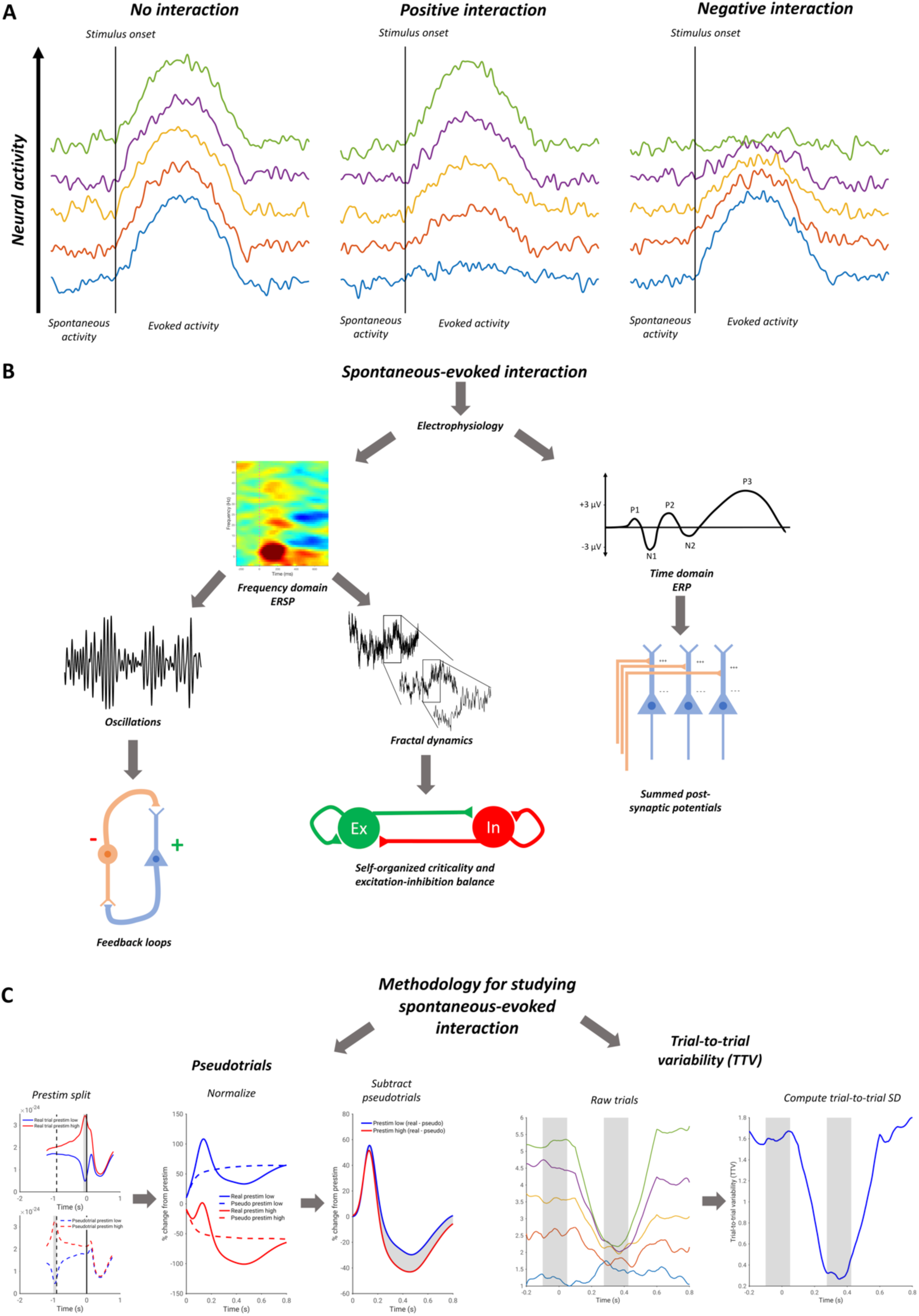
Schematic of the main aims and methods of the study. A) Schematic of different interaction schemes. With no spontaneous-evoked interaction, evoked amplitudes are identical regardless of the level of prestimulus spontaneous activity. In a positive interaction, higher prestimulus spontaneous activity leads to greater evoked amplitudes. In a negative interaction, higher prestimulus spontaneous activity leads to lower evoked amplitudes. B) Aims of the study. The study aims to assess which electrophysiological processes exhibit relationships between spontaneous prestimulus and evoked poststimulus activity. The study considers electrophysiological dynamics in the time and frequency domains, further classifying frequency domain event-related spectral perturbation (ERSP) as reflecting oscillations or scale-free (fractal) dynamics. Each of these electrophysiological parameters is associated with different physiological processes. C) Methodology for assessing spontaneous-evoked interaction. In the method of pseudotrials, trials and pseudotrials are split into prestimulus high and low conditions. They are then normalized relative to the mean prestimulus period, and the pseudotrial time courses are subtracted: any difference is indicative of a relationship between spontaneous and evoked activity. In the method of trial-to-trial variability, a negative interaction results in a reduction of the trial-to-trial standard deviation.

**Figure 2.**
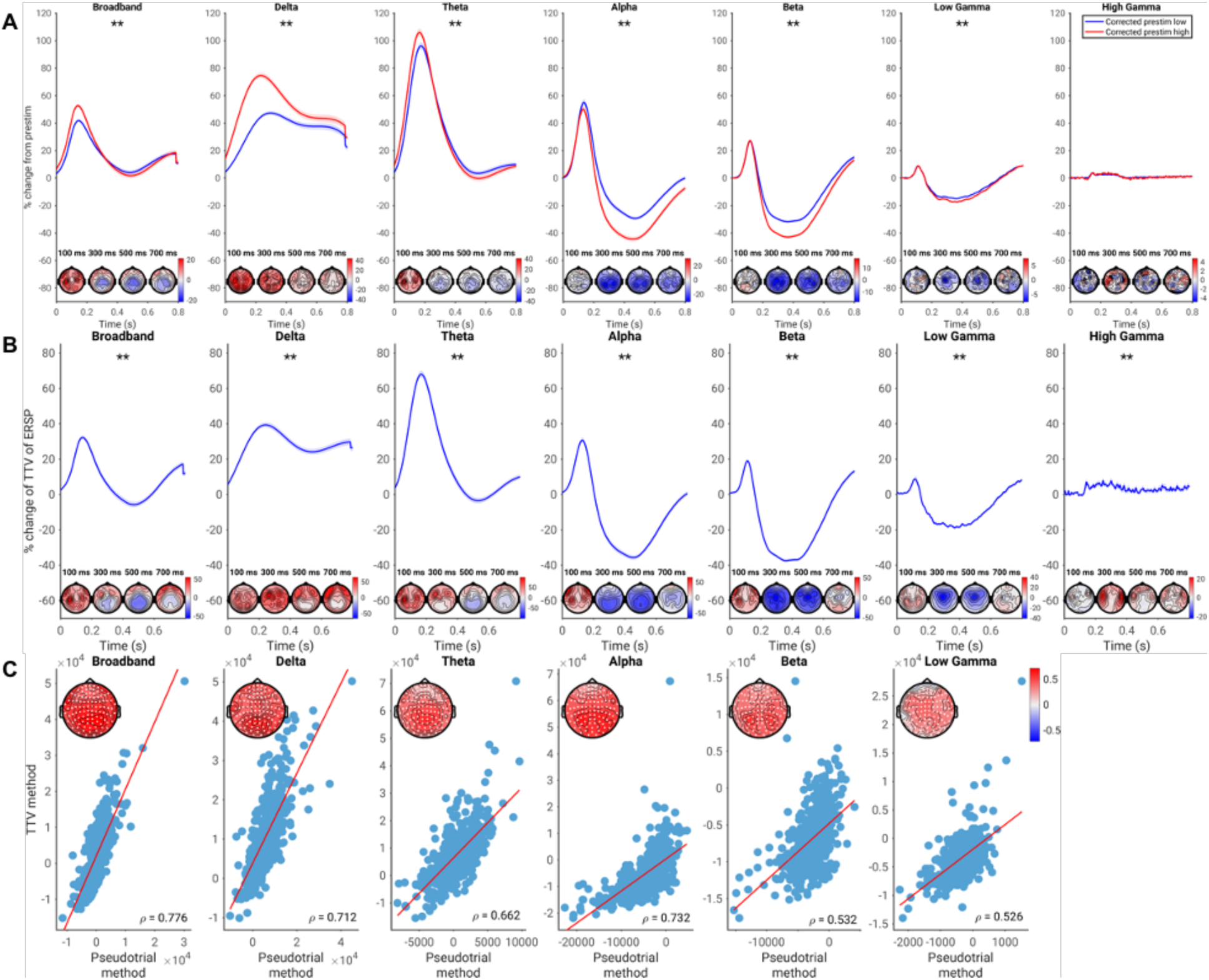
Spontaneous-evoked interaction in spectral power, assessed using the pseudotrial method (A) and TTV method (B). A) Pseudotrial-corrected time courses for high prestimulus (red) and low prestimulus (blue) conditions. Shaded area indicates standard error. Effect topographies are shown at 100 ms, 300 ms, 500 ms, and 700 ms. * = p < 0.05, ** = p < 0.01. B) Time course of TTV in each frequency band, expressed in terms of percent change from prestimulus levels. Asterisks and topographies as in A). C) Across-subject correlation of the magnitude of spontaneous-evoked interaction found using each method. Scatter plots show the correlation of the mean values across all electrodes, while the topoplots show the correlations at each electrode, with white dots indicating significance following the cluster test.

We observed both negative and positive interaction. Spontaneous-evoked interaction was negative (i.e., high-prestimulus trials lead to *lower* evoked activity compared with low-prestimulus trials) in beta and especially in alpha with a 17% difference at its maximum, peaking between 300 and 400ms post-stimulus. In contrast, positive interaction (i.e. high-prestimulus trials lead to *even higher* evoked activity compared with low-prestimulus trials) was found in the slower frequency bands of delta and theta, with a 33% maximum difference in delta peaking around between 150 and 250 ms post-stimulus.

We next confirmed the findings of an interaction between spontaneous and evoked spectral power using the method of trial-to-trial variability (TTV; figure 2b). We observed an early increase in TTV (between 100 and 200 ms) and subsequent TTV decrease (peaking around 400ms) in broadband (Figure 2b; p = 0.002). We then calculated TTV in different frequency bands. For the alpha (p = 0.002), beta (p = 0.002), and low gamma (p = 0.002) bands, we observed a highly significant decrease of the TTV (relative to the prestimulus period), which peaked between 400 and 500 ms.

In contrast, we observed initial increase in TTV in delta (p = 0.002) and theta (p = 0.002) bands, peaking between 150 and 250ms. A significant increase was also observed in high gamma (p = 0.002), though the magnitude of this effect was very small (around 5%). Note that as discussed in the methods, the presence of a TTV increase is not indicative of an interaction between spontaneous and evoked activity in and of itself (He 2013); however, we note that it does not contradict the results found using the pseudotrial method.

In order to validate that the two methods, i.e., pseudotrial and TTV, found similar magnitudes of spontaneous-evoked interaction, we calculated summary indices of spontaneous-evoked interaction by taking the signed area under the curve for each sensor and time point within the significant cluster (see Methods for details). We then correlated these summary indices across subjects at each electrode, correcting for multiple comparisons using a cluster-based permutation test (as no significant interaction was observed using the pseudotrial method in high gamma, we did not correlate this effect with the TTV method). We found that the two indices were significantly correlated in all frequency bands (Figure 2c; p = 0.002 in all cases, cluster-based test).

Using both pseudotrial-based and trial-to-trial variability-based methodologies, we show a positive interaction in the delta and theta bands, whereby high levels of prestimulus power lead to even greater ERSP increases. We also show a negative interaction in the alpha, beta, and low gamma bands, whereby high levels of prestimulus power lead to increased ERSP reduction post-stimulus. Results in the high gamma band were minimal in magnitude, and not significant using the pseudotrial method.

### Spontaneous-evoked interaction in the time domain – conflict between methodologies

We next tested for the possibility of a non-additive relationship between prestimulus and post-stimulus activity in the time domain, once again employing both the pseudotrial and TTV methods (Figure 3). Using the pseudotrial method, we observed a significant positive interaction in the time domain electrophysiological signal (Figure 3a; p = 0.002). Interestingly, this effect was largely constant throughout the post-stimulus period, rather than peaking and decreasing as with the power-based results. In contradiction to this finding, however, we observed (as seen in previous studies^59,66,67^) a reduction in TTV, occurring from 200 ms to 700 ms post-stimulus (p = 0.002), as well as an early increase in TTV (p = 0.006). This reduction in TTV, interpreted traditionally as in He (2013), should imply a negative interaction, in direct conflict with the results observed using the method of pseudotrials. These effects were significantly correlated (p = 0.002), indicating that less TTV reduction implies greater positive nonadditivity. However, the magnitude of the correlation was small (ρ of channel-average estimates = 0.218).

**Figure 3.**
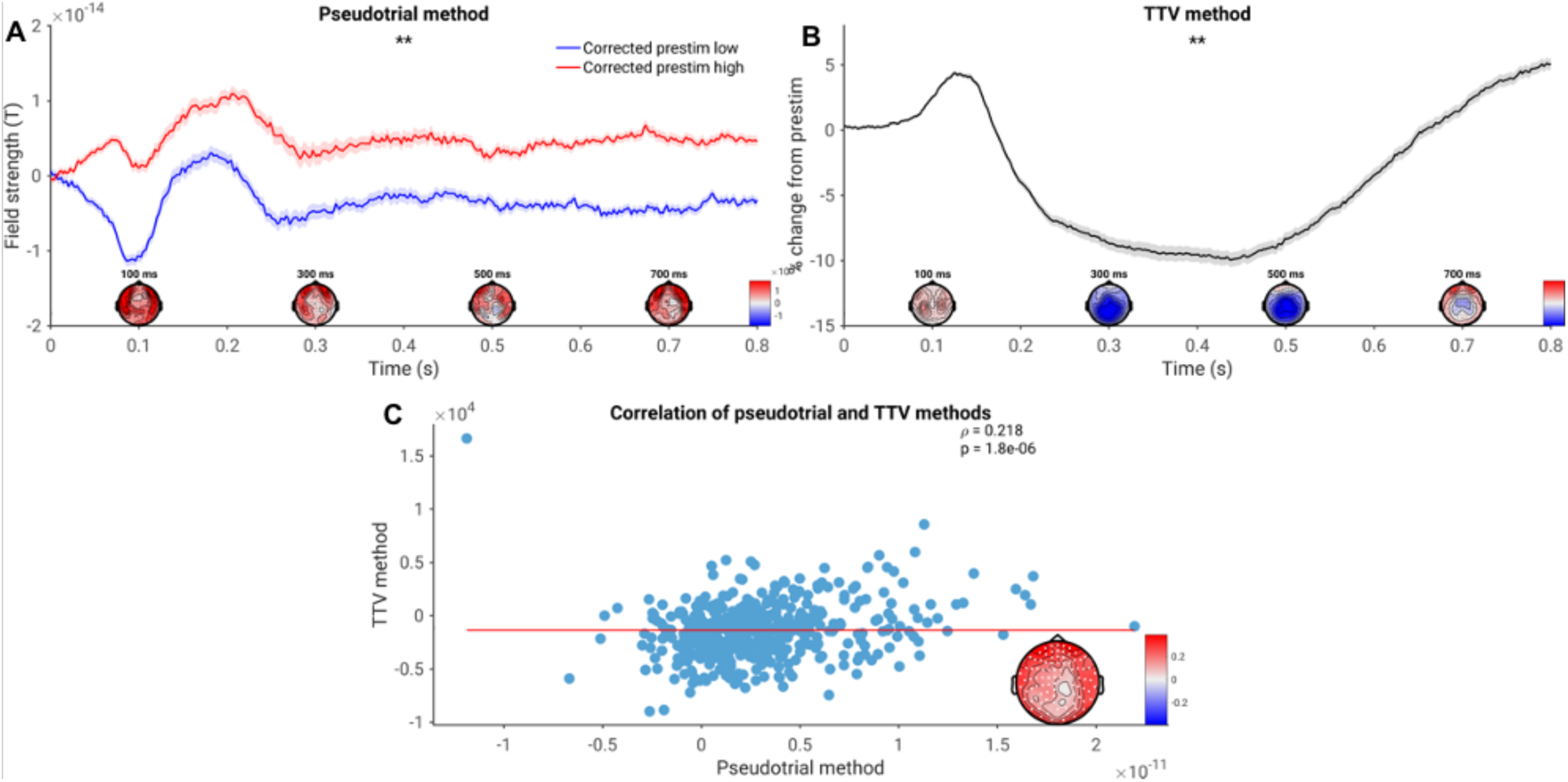
Time-domain spontaneous-evoked interaction of magnetic field strength, assessed using both methodologies. A) Pseudotrial-corrected time courses for high prestimulus (red) and low prestimulus (blue) conditions. * = p < 0.05, ** = p < 0.01. The shaded areas represent standard error. Effect topographies are shown at 100 ms, 300 ms, 500 ms, and 700 ms. B) Time course of trial-to-trial variability. Asterisks, shading, and topographies as in (A). C) Correlation of the magnitude of interaction assessed with each methodology. Mean values across sensors are plotted in the scatterplot, and the correlation coefficient at each electrode is plotted in the inset. White dots indicate sensors significant after cluster correction.

To examine possible sources of the discrepancy between the two methods, we conducted a simulation (Figure 4). As described in greater detail in the Materials and Methods, we simulated both a negative interaction with no change in oscillatory power, as well as an oscillatory power reduction with no interaction between prestimulus and poststimulus. We found that in both simulations, TTV decreased significantly (p = 0.002 in each case). However, only in the spontaneous-evoked interaction simulation did we observe a significant difference between prestim high and low using the pseudotrial method (p = 0.002). This suggests that the above findings of negative interaction using the TTV method may be confounded by the reduction of alpha power which also occurs following stimulus onset; for this reason, we view the results obtained using the method of pseudotrials as reflective of the genuine interaction pattern in the time-domain signal.

**Figure 4.**
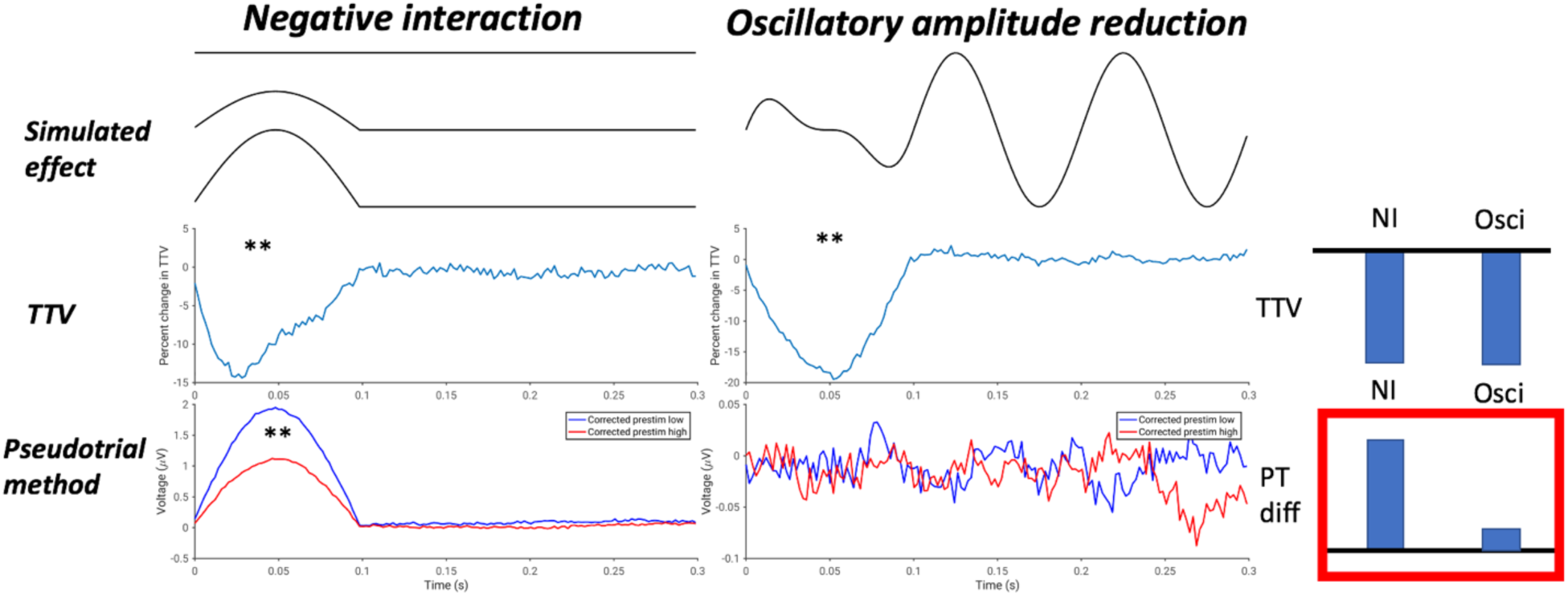
Results of simulation to disentangle methodological inconsistencies of figure 3. In each column, a schematic of the simulated effect is plotted (“simulated effect”), along with the normalized trial-to-trial SD (“TTV”), and the results following application of the method of pseudotrials (“pseudotrial method”). * = p < 0.05, ** = p < 0.01. Bars at the right show a schematic of the results: both simulations show a decrease in TTV, but only the true spontaneous-evoked interaction simulation shows a difference with the method of pseudotrials.

### Spontaneous-evoked interaction in the frequency domain - oscillatory and fractal components

The third aim of our study was to investigate whether the interaction between spontaneous and evoked activity differed between oscillatory and arrhythmic, scale-free processes. In order to examine more clearly the physiological substrates of prestimulus-dependent activity, we used the IRASA method^68^ to separate our data into oscillatory and scale-free (or “fractal”) components. We report abbreviated results of the application of this method to poststimulus activity in the supplementary materials (figure S1) as the IRASA method has never previously been applied to stimulus-locked activity; further detail on these results will be presented in a forthcoming publication.

Following the results of figure 4, we focused our analysis of spontaneous-evoked interaction in oscillatory and fractal power on the method of pseudotrials – results using the method of TTV are reported in the supplementary materials, and generally agree with the findings using pseudotrials (supplementary figure S8). Though it is more technically correct to study the fractal component in terms of its slope and intercept, in the following analyses we take the same frequency band view as in figure 2, in order to best compare fractal and oscillatory power with each other and examine each one’s contribution to the spontaneous-evoked interaction observed in the mixed power.

The findings using the method of pseudotrials show a positive interaction in delta and theta for the oscillatory component (Figure 6a; p = 0.002), as well as a negative interaction in alpha, beta, and low gamma (p = 0.002). Oscillatory high gamma displayed a positive interaction again (p = 0.002), but the magnitude of this effect was only a few percentage points. Similarly, the fractal component (Figure 6b) in the delta and theta bands displayed a positive interaction (p = 0.002), and beta and low gamma displayed a negative interaction (p = 0.002). However, it is noteworthy that in the fractal component, the alpha band displayed a small positive interaction, rather than a negative one as observed in the oscillatory component (p = 0.002). Once again, high gamma displayed a significant but small positive interaction (p = 0.002).

**Figure 5.**
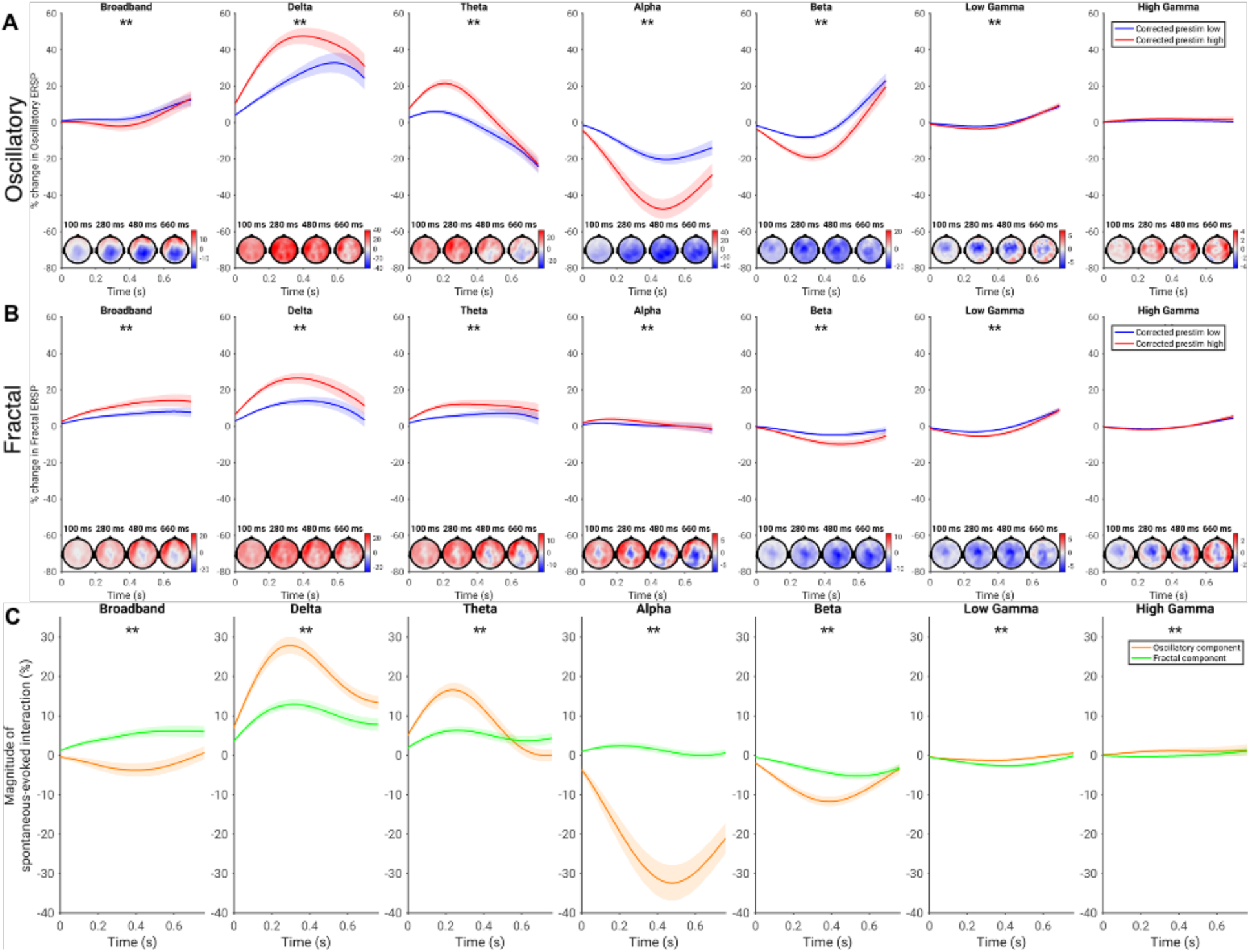
Interaction of spontaneous and evoked activity for oscillatory (A) and fractal (B) components of the power spectrum, assessed using the method of pseudotrials. A) Pseudotrial-corrected time courses for oscillatory power in high prestimulus (red) and low prestimulus (blue) conditions. Shaded area indicates standard error. Effect topographies are shown at 100 ms, 300 ms, 500 ms, and 700 ms. * = p < 0.05, ** = p < 0.01. B) As A), but for fractal power. C) Magnitude of the spontaneous-evoked relationship over time for oscillatory and fractal components. Asterisks indicate the significance of the difference between these two components.

**Figure 6.**
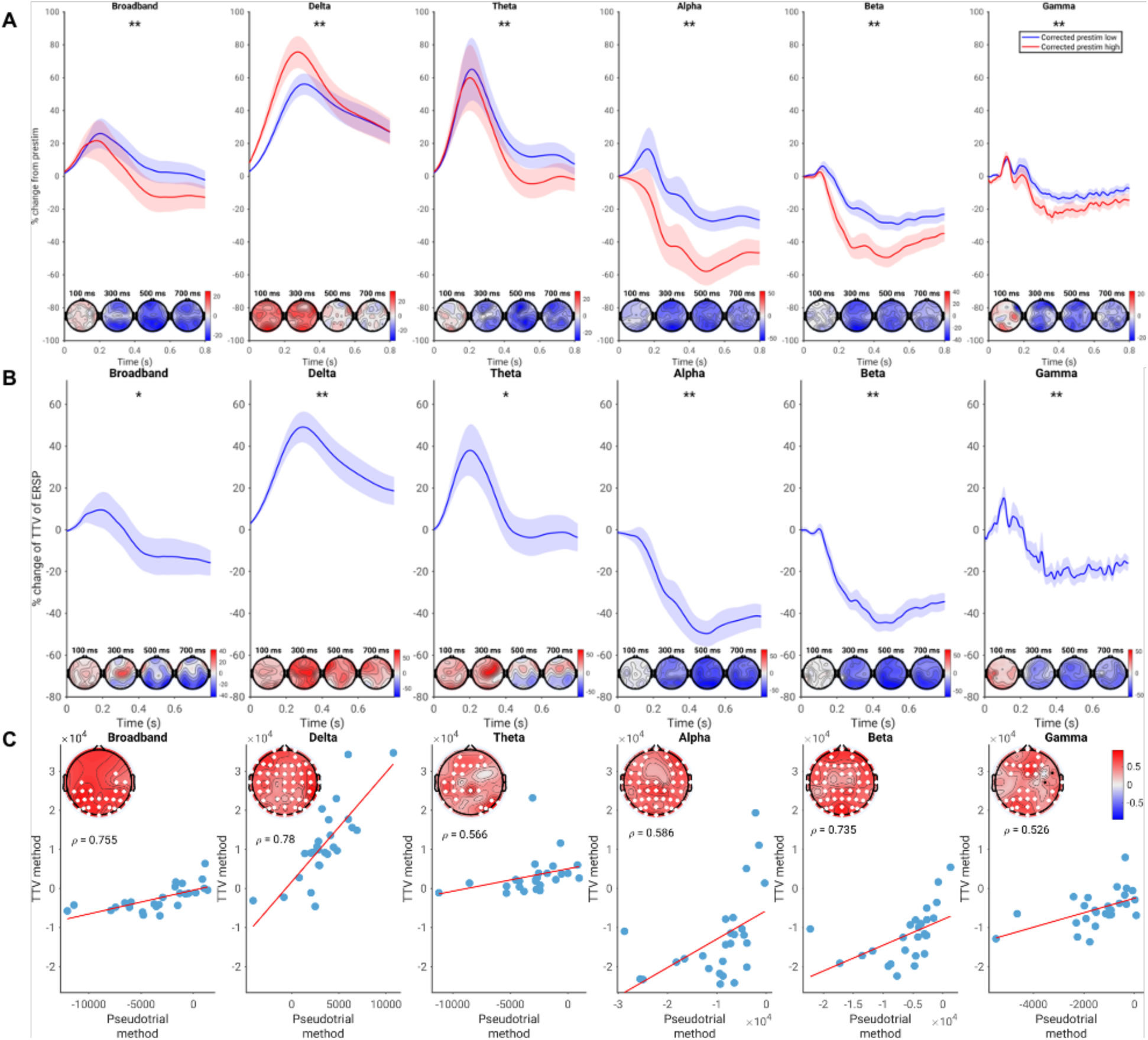
Spontaneous-evoked interaction of spectral power in the replication EEG dataset (equivalent to figure 2 for the main dataset). A) Pseudotrial-corrected time courses for high prestimulus (red) and low prestimulus (blue) conditions. Shaded area indicates standard error. Effect topographies are shown at 100 ms, 300 ms, 500 ms, and 700 ms. * = p < 0.05, ** = p < 0.01. B) Time course of TTV in each frequency band, expressed in terms of percent change from prestimulus levels. Asterisks and topographies as in A). C) Across-subject correlation of the magnitude of interaction found using each method. Scatter plots show the correlation of the mean values across all electrodes, while the topoplots show the correlations at each electrode, with white dots indicating significance following the cluster test.

Comparing the magnitudes of the effects, we see that in most cases the oscillatory component displayed a much stronger interaction, particularly in the delta, theta, alpha, and beta bands, as well as high gamma (Figure 6c; p = 0.002). The fractal component displayed a slightly stronger interaction in the low gamma band, but the effects were again small compared with the other bands (p = 0.002). Together, our results suggest distinct contributions of oscillatory and arrhythmic/fractal components to positive and negative interaction schemes of spontaneous and evoked activity.

### Replication of spontaneous-evoked interaction in an independent EEG dataset

To ensure robustness of the findings, we replicated our procedure in an independent EEG dataset (figures 7 and 8). This dataset consisted of a more cognitive paradigm, a moral decision-making task previously described in Wolff et al.^59^ Using the method of pseudotrials, we generally found the same pattern of spontaneous-evoked interaction as in figure 2, with delta displaying a positive interaction and alpha, beta, and gamma showing negative interaction (Figure 7a; p = 0.002). In contrast, however, we observed a negative interaction in the theta band in the EEG data (p = 0.002). We submit that this difference in theta may be due to the band being a transition between the positive interaction regime in delta and the negative one in alpha and beta. We can see in the EEG data, the delta ERSP is smaller relative to the alpha ERSP than in the MEG data; regardless of whether this is due to task effects or the difference in imaging modality, this difference in relative contribution may explain the different interaction scheme in theta.

**Figure 7.**
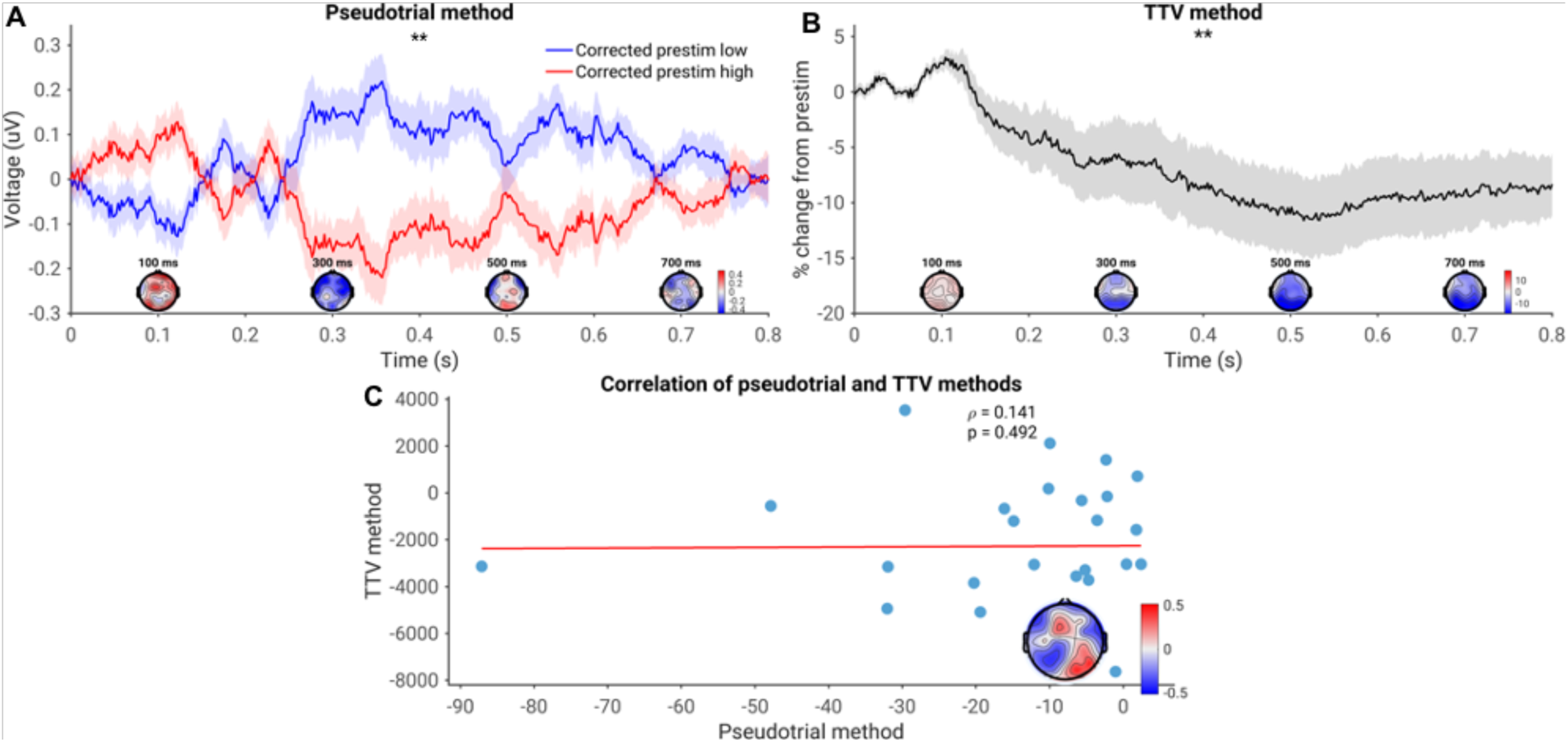
Time-domain spontaneous-evoked interaction in EEG voltage for the replication EEG dataset (equivalent to figure 3 in the main dataset). A) Pseudotrial-corrected time courses for high prestimulus (red) and low prestimulus (blue) conditions. * = p < 0.05, ** = p < 0.01. The shaded areas represent standard error. Effect topographies are shown at 100 ms, 300 ms, 500 ms, and 700 ms. B) Time course of trial-to-trial variability. Asterisks and topographies as in (A). C) Correlation of the magnitude of spontaneous-evoked interaction assessed with each methodology. Mean values across sensors are plotted in the scatterplot, and the correlation coefficient at each electrode is plotted in the inset. White dots indicate sensors significant after cluster correction.

The results using the method of trial-to-trial variability largely confirm the results obtained using the method of pseudotrials (Figure 7b; p = 0.036 for broadband, p = 0.024 for theta, p = 0.002 for each other band). The only exception is that the TTV method does not indicate a negative interaction in the theta band – however, as mentioned by He (2013) and in the introduction, this does not mean that such an interaction is definitively not present, only that the variability of the spontaneous and evoked activity is sufficient to overwhelm the term related to their correlation in this case. For all bands, the magnitudes of interaction calculated with each method were correlated (p = 0.002 for each band).

We also assessed spontaneous-evoked interaction in the time domain in EEG data. Similar to the MEG data, we observed a decrease in TTV over much of the poststimulus period (Figure 8b; *p* = 0.002). However, in contrast to the MEG findings, we observed a negative interaction in the EEG data (Figure 8a; p = 0.002). These findings were uncorrelated (Figure 8c; p = 0.246). See the discussion for further treatment of this difference from the MEG data.

### Control analyses - simulations

We additionally ensured that the length of the prestimulus period did not affect the findings. In addition to the presently used 100 millisecond window, a 50-ms prestimulus period (Supplementary figures S4-S5) and a 200-ms prestimulus period (Supplementary figures S6-S7) were used. These values had little discernable effect on the main results.

In order to distinguish true interaction between spontaneous and evoked activity from potential effects due to trial-by-trial variations in noise, we tested for spontaneous-evoked interaction in two simulations: one in which a purely additive (additive simulation; supplementary figure S2) relationship was present, and another in which a non-additive (negative) interaction occurred (non-additive simulation; supplementary figure S3). In the additive simulation, no evidence of spontaneous-evoked interaction was observed using the pseudotrial method. However, several significant decreases of TTV were observed, as well as increases in different noise conditions. In contrast, in the non-additive simulation, we observed consistent negative interaction using the method of pseudotrials and the method of TTV in the low-noise conditions. This confirms that the method of pseudotrials is capable of detecting true spontaneous-evoked interactions given a sufficient signal-to-noise ratio, and does not suffer from a high degree of false positives due to time-varying noise. The method of TTV also appears capable of detecting a spontaneous-evoked interaction, but in certain noise regimes it can be subject to false positives.

We also attempted to control for the potential inclusion of part of the post-stimulus response in the pre-stimulus period. This could occur for a variety of reasons, including high-pass filtering in the case of the time-domain signal (Supplementary figure S9), and because of the inherent trade-off between time and frequency resolution in the case of frequency-domain signals (Supplementary figure S10). We found that while these factors may have resulted in overestimation of the magnitude of positive interaction in our data, positive interaction remained significant when controlling for them.

## Discussion

The goal of our study was to examine the interaction between spontaneous and evoked activity and its neuronal mechanisms, using the general framework of negative and positive interaction in electrophysiology. We showed strong relationship of spontaneous activity in the prestimulus period with evoked activity in the poststimulus period; this was observable in both frequency-domain and time-domain analyses. The observed correlation was negative in the alpha and beta bands (such that high alpha/beta amplitude pre-stimulus leads to greater stimulus-evoked desynchronization), and positive in the delta band (such that high delta amplitude pre-stimulus leads to greater stimulus evoked synchronization). Moreover, we demonstrate that this effect is primarily found in band-limited oscillatory dynamics, rather than arrhythmic, scale-free dynamics. Finally, we also observed spontaneous-evoked interaction in the time domain, with a positive interaction observed in MEG in the multisensory task, and a negative interaction in the EEG moral reasoning task.

These results bridge the gap between spontaneous and evoked activity by showing that trial-by-trial fluctuations in pre-stimulus activity influence fluctuations in post-stimulus neural responses to stimuli in predictable ways. Our results extend recent findings on relating prestimulus spectral power to ERP components^69,70^, and complement cellular and modelling research investigating this question^71–74^. This research contributes to a more complete understanding of how fluctuations in spontaneous activity mediate variability in behavioral and cognitive features like perception, self, attention, and consciousness.

### Spontaneous-evoked interaction schemes - positive and negative interaction in different frequency bands

We demonstrate a non-additive relationship between prestimulus and poststimulus activity in multiple electrophysiological mechanisms. For spectral power, we showed similar spontaneous-evoked interaction patterns in different paradigms and modalities, i.e., sensory in MEG and cognitive in EEG, as well as using different methods (TTV^43^ and pseudotrial^44,59^). This suggests that the close relationship between spontaneous and evoked activity in spectral power is a robust phenomenon that holds across different tasks and methods. Most interestingly, we demonstrated two different interaction schemes holding in the spectral power of different frequency bands. Negative interaction was predominant in alpha in the later post-stimulus periods (around 400ms). In contrast, positive interaction was observed in the delta band, occurring earlier around 200ms; such an interaction scheme has not yet been reported in the literature.

The existence of a negative interaction in specifically alpha is noteworthy given prior data showing the impact of pre-stimulus alpha on post-stimulus perception^27,28,75^, self^30^, and conscious awareness of stimuli^32,75,76^. Alpha has traditionally been regarded as an inhibitory process, with alpha desynchronization reflecting release from inhibition; according to Klimesch^77^, this reflects controlled access to the “knowledge system”. Furthermore, prestimulus or anticipatory alpha desynchronization has been associated with attention allocation and better subsequent performance on perceptual discrimination tasks^78,79^. Therefore, we suggest that the observed interaction in alpha may reflect the phenomenon of pre-allocated attention and awareness in the subject leading to better behavioural performance. Trials with low prestimulus alpha do not require release from cortical inhibition, as inhibition is already low and anticipatory attention has been allocated, so alpha desynchronization is less prominent. In contrast, for trials where prestimulus alpha power is high, more desynchronization is needed in order to respond effectively. As such, the negative interaction we observed may reflect the dependence of the post-stimulus alpha response and the associated cognitive processes on pre-stimulus alpha amplitude.

Additionally, we observed positive interaction in the delta band. Delta-band activity has been shown to mediate arousal and attention in the waking state^80–82^, including mediating the p300 event-related potential^83,84^. It is also known as an electrophysiological correlate of activity in the default-mode network^85^, and as a modular of motivation, arousal and homeostasis^186^. Though our findings leave open the exact function of positive interaction in delta, it may provide a mechanism linking motivation and arousal with responses to salient stimuli. However, we note that the positive interaction in delta may be overestimated due to the temporal imprecision associated with estimating slow-frequency activity (see Supplementary figure S10), and urge caution in the interpretation of these results.

Our finding of a non-additive interaction between pre- and post-stimulus activity carries methodological implications for the analysis of evoked activity, where it is tacitly assumed that stimulus-evoked responses are superimposed additively upon spontaneous fluctuations. Particularly in electrophysiology, trial-based paradigms are the dominant way of conducting task-related neuroscience research^87^. Our results suggest that the assumption of trial independence that underlies these paradigms is problematic – as a result, these studies may miss important data related to the dependence of the evoked response on pre-stimulus and ongoing activity. Our data, emphasizing the impact of continuous fluctuations in spontaneous activity on transient stimulus-evoked responses, provide support for recent attempts to develop analysis strategies for continuous paradigms which do not require the averaging of multiple trials^87^.

### Electrophysiology of spontaneous-evoked interaction – time vs. frequency domains, oscillatory vs. fractal dynamics

We observed a consistent interaction between spontaneous and evoked activity in spectral power, i.e., ERSP. The spontaneous-evoked interaction in spectral power was observed for both methods, i.e., TTV and pseudotrials, as well as in both modalities, i.e., MEG and EEG. Given that ERSP is believed to reflect changes in interactions between multiple neurons in thalamo-cortical and cortico-cortical feedback loops^49^, future modelling studies should investigate how the properties of these systems and networks can produce state-dependent effects in the ways observed here. These findings suggest that pre-post-stimulus interaction is a feature of complex inter-cellular and inter-regional feedback or recurrent mechanisms.

Using the method of pseudotrials, we also observed a spontaneous-evoked interaction in the time-domain electrophysiological signal in both modalities. However, we observed differences in the spontaneous-evoked interaction of the time-domain signal between the MEG and EEG datasets. This may reflect a difference between the two modalities, as MEG and EEG are sensitive to different cortical sources^88^. Alternatively, it could be that the task (simple sensory vs. complex cognitive) may have an effect on the interaction scheme exhibited by the time-domain signal. However, because of the methodological issues pointed out in figure 4 and figure S9, more research is needed to investigate spontaneous-evoked interaction in time-domain electrophysiological signals and the conditions that affect it.

Finally, we observed a larger influence of spontaneous activity on evoked responses in the oscillatory rather than the scale-free component of the power spectrum. This complements recent observations of the differential relationships of oscillatory and scale-free dynamics to the fMRI signal^61^, differential contributions to cognitive processing speed^89^, and differential modulation by psychedelic drugs^90^. It also reinforces the central role of cortical oscillations in stimulus response, and may point to a more “background” dynamic role for scale-free activity in the brain, in which scale-free activity influences cognitive processing without being directly involved in it^91^.

### Psychiatric applications of spontaneous-evoked interaction

As a mechanistic link between prestimulus activity and poststimulus activity, spontaneous-evoked interaction also has the potential to connect a number of disparate observations in psychiatric disorders. It is often observed that psychiatric patients have abnormalities in the level of both spontaneous and evoked spectral power^92,93^. For example, Keehn et al.^94^ observed lower spontaneous alpha power in autism, which correlated with reduced alpha ERSP response. Similarly, in patients with alcoholism, studies have found reduced spontaneous delta amplitude^95,96^, as well as reduced delta ERSP^97,98^.

Our data suggest that both of these sets of observations can be explained parsimoniously within the negative/positive interaction framework advanced in our study. For example, given the negative interaction we observed in alpha, lower spontaneous pre-stimulus alpha power should lead to less post-stimulus alpha desynchronization – exactly the finding observed by Keehn et al. in autism. Similarly, given the positive interaction observed in delta ERSP, lower levels of spontaneous pre-stimulus delta activity should lead to a reduction in the poststimulus delta ERSP response – again, the exact finding observed in alcoholic patients. Our results provide a link between the evoked and spontaneous activity abnormalities in the psychiatric disorders described above. As such, the interaction between spontaneous and evoked activity is a promising candidate for future research in psychiatric disorders, and possible mechanism by which therapies which alter spontaneous activity might consequently modulate task-evoked activity and behaviour.

### Limitations

As discussed above, a central limitation of our study is the methodological difficulty of assessing interaction between spontaneous and evoked activity when activity in the post-stimulus period reflects a mixture of the two. Two main issues emerged in our study: the failure of the TTV method in time-domain electrophysiological data, and the confounding of positive interaction estimates by temporal imprecision. Because it relies on the law of total variance, the method of TTV assumes stationarity of spontaneous and evoked activity (i.e. that their variances are approximately constant over time); this assumption holds in fMRI^99^, but is broken in EEG and MEG^100^. As shown in figure 4, TTV decreases in the raw electrophysiological signal can be induced by desynchronization of oscillations. The method of pseudotrials appears still to be effective here; however, in the case of positive interaction, the method of pseudotrials can be affected by any mechanism which “smears” poststimulus activity back into the prestimulus period, such as highpass filtering in the time domain (supplementary figure S9), or the inherent imprecision in estimating low-frequency power (supplementary figure S10). While all our results remain significant while considering these effects, this issue does appear to result in overestimation of the magnitude of positive interaction, and as such more research is needed to confirm the positive interactions seen in our data. In particular, methods which focus on inter-subject consistency of responses in order to isolate task-evoked activity (such as the one applied by Lynch et al.^101^) may be useful in these cases, but they have not yet been developed for the purpose of relating spontaneous and evoked activity.

We were not able to determine a particular mechanism which explained the interaction patterns we observed. In a preliminary investigation, we found no influence of phase coherence or phase-amplitude coupling (supplementary figure S11) on the negative interaction scheme observed in our study. However, we did not investigate these possibilities in detail; further empirical and modelling work is necessary to determine more precisely why and how spontaneous and evoked activity interact in the ways we observed.

## Conclusion

In this paper, we investigated the interaction between prestimulus and poststimulus activity and its manifestation in multiple electrophysiological processes. Using both MEG and EEG and applying robust analysis methods, we observed, as hypothesized, that spontaneous activity in the pre-stimulus period exerted an influence on evoked activity. Positive interaction was found in evoked delta power, while negative interaction occurred in the alpha and beta bands. Both forms of spontaneous-evoked interaction were found robustly in the dynamics of spectral power, and predominantly in oscillatory rather than arrhythmic/scale-free dynamics. Both forms of interaction could also be observed in the time-domain electrophysiological signal, with differences between the MEG and EEG datasets.

This work carries major methodological implications for our understanding of stimulus-induced or task-evoked activity by unraveling, in part, the mechanisms by which it is related to spontaneous neural activity. Even more importantly, these findings provide insight into the neuronal mechanisms of spontaneous-evoked interaction which necessarily underlie the well-documented effects of spontaneous activity on perception, cognition, and consciousness.

## Materials and methods

### Methodologies for assessing correlations between spontaneous and evoked activity

fMRI studies investigating pre-post-stimulus interaction have employed two distinct methods: trial-to-trial variability (TTV)^43^ and pseudotrials^44^. The method of *trial-to-trial variability* (TTV) makes use of the law of total variance in assessing a correlative relationship between spontaneous activity (X) and evoked activity (Y):

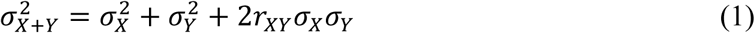

A putative interaction is represented by the correlation *r*_*XY*_. Since variances are always positive, the only circumstance in which one could observe a reduction in trial-to-trial variability is if this coefficient is negative – hence, a reduction in TTV implies a negative correlation between spontaneous and evoked activity. However, an increase in TTV could be produced by any of the three models. It was found that the trial-to-trial variability of the fMRI response decreased following stimulus onset, a pattern which could only be explained assuming a negative interaction model. This phenomenon has frequently been associated with attractor models of the brain^102–104^.

More recently, a more direct method has emerged to assess the presence of spontaneous-evoked interaction in fMRI^44^. The method of *pseudotrials* directly assesses the spontaneous-evoked interaction by splitting trials at the median based on pre-stimulus activity and comparing the average time course for the trials with low prestimulus activity and the trials with high prestimulus activity. However, such an approach can be confounded by regression to the mean^60^ – trials which are selected based on having high prestimulus activity may naturally return to a more average value, simply because they are selected as having “above average” values in the first place, or by other dynamical features of the spontaneous activity.

To correct for this, Huang et al.^44^ applied the same procedure to “pseudotrials” (segments of the task recording where no stimulus is present) and subtract the pseudotrial time courses from the real trial time courses. In this way, a direct estimate of the influence of spontaneous activity on evoked activity is made which corrects for regression to the mean, and can theoretically detect a positive interaction and distinguish it from a negative one (which cannot be accomplished with the method of trial-to-trial variability). Using this method, a negative interaction was observed in fMRI, which correlated with the same interaction assessed with the TTV method^44^.

### Datasets and experimental designs

We used two previously published datasets to investigate our hypotheses, in two different imaging modalities. As our main dataset, we used 474 subjects from the Cambridge Centre for Ageing and Neuroscience (CamCAN) MEG dataset^57,58^ (available at http://www.mrc-cbu.cam.ac.uk/datasets/camcan/). Data were recorded using a Vectorview 306-channel MEG system (Elekta Neuromag, Helsinki) in a light magnetically-shielded room. The task associated with this dataset was a simple sensorimotor task, in which participants were presented with a multimodal auditory (300 ms tone) and visual (34 ms checkerboard pattern) stimulus. 120 trials were presented, and the inter-trial intervals (ITIs) were jittered between 2 and 26 seconds. The auditory and visual stimuli were presented simultaneously, and participants were required to respond with a button press once they perceived the stimulus. Eight trials were also included of unimodal auditory or visual stimuli (four trials each) – these were also included in the analysis. Details of the task can be found in ^58^.

To replicate the findings, we used data from a moral decision-making paradigm, previously published in^59^. Participants in this task were presented with a moral dilemma, in which participants would push a set of bystanders in front of a trolley in order to save another group from death (Philippa Foot’s “Footbridge dilemma”). Participants were asked to respond either yes or no to the ratio of those killed to saved (in visual stimulus) by pushing a button. An additional externally-guided condition required participants simply to compare the number of people on the right side of the screen with the left. For the purposes of our study, we considered both conditions together. A total of 420 trials were included in the analysis, and the ITI was jittered between five and six seconds. Further details can be found in Wolff et al.^59^

### Data preprocessing

At the time of analysis, fully preprocessed data were not available from the CamCAN repository. As such, we applied standard preprocessing steps in Fieldtrip^105^ and MNE^106^ to further clean the data. Our starting point for preprocessing was the MaxFiltered data provided in the CamCAN data release (see^57^ for details of the MaxFilter steps). For ease of preprocessing and analysis, gradiometer channels were removed, and only magnetometers were analyzed. Data were first downsampled to 500 Hz, then bandpass filtered from 1 to 200 Hz with a fourth-order Butterworth filter. In order to remove high-amplitude transient artefacts which could bias independent component analysis (ICA) decomposition, Autoreject^107^ was applied to find and label data epochs with artefacts. Following the methods of the Human Connectome Project^108^, 20 ICA iterations were then performed, with artefactual epochs from the previous step excluded from the ICA training. Artefactual components were automatically labelled using in-house modifications of scripts from the Human Connectome Project’s megconnnectome software^108^ and removed from the original (pre-Autoreject) data. Data were then epoched from 2 seconds prior to stimulus onset to 1.5 seconds poststimulus, and Autoreject was run a second time to repair any trials with artefacts remaining.

Data from the moral decision-making paradigm were preprocessed in EEGLAB^109^. First, data were downsampled to 500 Hz and bandpass filtered from 1 to 50 Hz. High-amplitude artifacts were removed prior to ICA using Artefact Subspace Reconstruction^110^. Data were then epoched from 3 seconds prior to stimulus onset to 2 seconds poststimulus. ICA was then run on the data, and bad components were identified using an automated algorithm^111^. For details of the data collection, see Wolff et al.^59^

### Definition of real trials and pseudotrials

For each dataset, he prestimulus period for real trials was taken as the interval from 100 milliseconds before stimulus onset to stimulus onset, with the real trial poststimulus period falling from 0 to 800 milliseconds. Similarly, “pseudotrials” were defined as the period from 900 to 100 milliseconds pre-stimulus, with the period from 1000 to 900 milliseconds pre-stimulus serving as the “prestimulus” period for that pseudotrial. We tried several different lengths of the prestimulus interval, to ensure that this choice had minimal impact on the results. These findings are reported in the supplementary materials (figures S4-S7).

### Estimation of negative and positive interaction

We used both the method of pseudotrials^44^ and the method of trial-to-trial variability^43^ to assess the presence of a non-additive relationship between spontaneous and evoked activity in various signals. We averaged the amplitude across time points in the prestimulus window to get a single value characterizing prestimulus amplitude for each subject, channel, and trial. For each subject, we then split trials based on the median into high or low prestimulus amplitude. This procedure was carried out separately for pseudotrials and real trials. For spectral power, pseudotrial and real trial time courses were then normalized by the average prestimulus value over all trials and expressed in terms of percent change from this value. This normalization was not done for time-domain signals, as the average prestimulus value was approximately zero.

We then computed the “corrected” time courses by subtracting the normalized real trial time courses from the normalized pseudotrial time courses (see^44^). This “corrected” time series is now controlled for regression to the mean, as this natural difference between prestim high and low is captured by the pseudotrial dynamics and removed from the data. We then tested for whether the corrected time series for high and low prestimulus amplitude were significantly different from each other using the cluster-based procedure described below.

For the method of trial-to-trial variability, we calculated the standard deviation across trials of the signal of interest. This time course was normalized to its mean value in the 100 millisecond prestimulus period and expressed in percent change from this value. A decrease in TTV indicates a negative interaction, while an increase in TTV cannot distinguish between additivity and positive interaction^43^.

### Time-frequency decomposition and definition of frequency bands

Time-frequency decomposition was carried out using the wavelet transform, as implemented in Fieldtrip. Three cycles of the default Morlet wavelet were used to estimate 50 logarithmically-spaced frequencies from 2 to 200 Hz (or from 2 to 50 Hz in the EEG case). The delta band was defined as 2-4 Hz, low gamma as 30-100 Hz (or 30-50 Hz in the case of the EEG dataset), and high gamma as 100-200 Hz. Since alpha peak frequency and peak width varies significantly between individuals, we followed the recommendations of Klimesch^112^ and defined the alpha, theta, and beta bands individually for each subject. We estimated the alpha peak frequency and width using the toolbox developed by^113^. The alpha band was then defined according to this width: the theta band was subsequently defined as 4 Hz to the lower bound of the alpha band, and beta was defined as the upper bound of the alpha band to 30 Hz. For subjects where no alpha peak was found, these bands were defined in the standard way (theta as 4-8 Hz, alpha as 8-13 Hz, and beta as 13-30 Hz).

### Separation of oscillatory and fractal components of ERSP

To distinguish oscillatory and fractal processes in the spectral power response, we used the Irregular Resampling for Auto-Spectral Analysis (IRASA) method, described in^68^. In brief, this method involves resampling the signal using a variety of non-integer resampling factors and taking the median of the resulting power spectra to represent the fractal component. For this analysis, we used a subset of subjects from the main CamCAN dataset (n = 49), due to the extreme computational intensiveness of the time resolved IRASA procedure.

Following a previous publication^90^, we used resampling factors ranging from 1.1 to 2.9 in steps of 0.05, excluding the factor 2. To assess stimulus-related changes in oscillatory and fractal ERSP, we computed this method in a sliding window of size 1.5 seconds and step size 20 ms. This allowed us to estimate frequencies ranging from 2 to 125 Hz. The frequency bands for this process were the same as for the wavelet-based ERSP, but since only frequencies up to 125 Hz could be resolved using the IRASA method, low gamma was defined as 30 to 50 Hz and high gamma as 50 to 125 Hz.

Occasionally, the IRASA method returns negative values of oscillatory power because oscillatory power is computed as the difference of the mixed power spectrum and the fractal component. Negative values of oscillatory power were treated as missing data in this study.

### Statistical analysis

In order to achieve good statistical power and flexibility without a strong *a-priori* hypothesis as to the latency or topography of the pre-poststimulus interaction, we opted to used cluster-based permutation tests^114^. Our time window of interest was defined as the time between 0 and 800 milliseconds poststimulus. We tested for significant difference between the corrected prestim low and high time courses using a Wilcoxon signed-rank test at each time point and each sensor, using a cluster-based permutation test with 1000 permutations to correct for multiple comparisons. For the TTV-based method, we used a signed-rank test against zero, similarly using a cluster-based permutation test to address the multiple comparison problem. Both pseudotrial-based and TTV-based spontaneous-evoked interaction indices were then calculated only for the sensors and time points which formed part of a significant cluster. As rightly pointed out by Sassenhagen and Draschkow^115^, the permutation test does not establish the significance of the latency or topography, per se – we considered it here merely as a useful prior for calculating a summary index of the magnitude of spontaneous-evoked interaction. These indices were then correlated across subjects using Spearman’s rank correlation.

### Simulation of the discrepancy between pseudotrial method and TTV method

It has recently been observed that in EEG, TTV and spectral power are closely related^116^. This could present a confounding factor for the assessment of pre-post-stimulus interaction in the time-domain signal. To assess this possibility, we carried out two simulations: true spontaneous-evoked interaction in the time-domain signal, and oscillatory power reduction with no additional time-domain stimulus response. Forty-eight subjects and 128 4-second long trials were simulated using an in-house modification of Fieldtrip’s ft_freqsimulation function. In each trial, electrophysiological data were simulated as a summation of 1/f^β^ noise (β randomly chosen between 0.5 and 1.5) and an alpha oscillation at 10 Hz. We modelled the amplitude of this oscillation itself as a 1/f^ß^ process lowpass filtered at 1 Hz. In the spontaneous-evoked interaction simulation, a stimulus related increase in the form of one lobe of a sine function was added to the signal; the magnitude of this increase was varied according to the value of the prestimulus voltage. In the oscillatory power reduction simulation, the amplitude of the 10 Hz oscillation was reduced over the same time period, again with the response taking the form of one lobe of a sine function. We then tested for spontaneous-evoked interaction in each noise regime using the method of pseudotrials and the method of TTV, as described above. As in our real data, we used a cluster-based permutation test, clustering only across the time dimension as only one channel was simulated.

### Simulation of additive and non-additive models accounting for trial-varying signal-to-noise ratio

A correlation between spontaneous and evoked activity could conceivably be observed due to trial-varying signal-to-noise ratio. If SNR is high on some trials, one may observe an association between higher prestimulus power and a greater ERSP response simply due to the ERSP estimation being less corrupted by noise in these trials. To test for this possibility, we conducted simulation experiments in which we systematically varied the signal-to-noise ratio across trials (Supplementary figures S2 and S3). We varied both the base SNR and the across-trial variability of the noise across four orders of magnitude (from 1/64 to 64 in each case). Under each noise regime, we tested two models: one, in which the evoked response and the ongoing dynamics were independent (negative control; supplementary figure S2), and another in which the evoked response and ongoing dynamics were correlated (positive control; supplementary figure S3). The simulation was carried out as described above for the oscillatory power reduction simulation (though the ERSP response was extended to better match the observed temporal characteristics of the alpha response). The magnitude of this decrease was either scaled by the spontaneous amplitude in the 100 millisecond prestimulus period (positive control; spontaneous-evoked interaction) or varied randomly (negative control; additive interaction). Statistical tests were carried out as described above.

## Acknowledgements

Data collection and sharing for this project was provided by the Cambridge Centre for Ageing and Neuroscience (CamCAN). CamCAN funding was provided by the UK Biotechnology and Biological Sciences Research Council (grant number BB/H008217/1), together with support from the UK Medical Research Council and University of Cambridge, UK. This project has received funding from the European Union’s Horizon 2020 Framework Programme for Research and Innovation under the Specific Grant Agreement No. 785907 (Human Brain Project SGA2). GN is grateful for funding provided by UMRF, uOBMRI, CIHR, and PSI.

