## Supplementary materials for "Bridging the gap – Spontaneous fluctuations shape stimulus-evoked spectral power"

*Spectral power response of fractal and oscillatory components*

While the IRASA method has been applied in the context of continuous, long-term EEG and MEG recordings[1,2], it has not been applied in a time-resolved manner to stimulus related activity. Here we report (in brief) the stimulus-related changes of oscillatory power and fractal power in each frequency band; these results will be reported in more detail in a forthcoming publication. We note also that the fractal component is not traditionally divided into frequency bands – indeed, this contradicts the nature of the fractal component as “scale-free”, broadband activity. We follow here the same approach as in the main text, which considered the fractal component in a band-limited setting in order to best contrast it with the oscillatory component and make use of the methods available for assessing spontaneous-evoked interaction. Results considering the fractal component from a parametric view (i.e. considering its slope and intercept) will also be reported in a forthcoming publication.

We observed a significant spectral power change in each frequency band, in both the fractal and oscillatory components ($p = 0.002$ in each band). The ERSP response of the oscillatory component was familiar; it shows an increase in delta- and theta-band activity which peaks early in the post-stimulus period (250 ms), and a decrease in theta-, alpha-, beta-, and low gamma-band activity which bottoms out later in the post-stimulus period (400-600 ms). High gamma band activity showed a very small post-stimulus increase. The ERSP response of the fractal component was in general similar; slow-frequency power increased, and fast-frequency power decreased. Delta and theta power increased, peaking slightly between 200 and 400 ms post-stimulus, but remaining high for the rest of the post-stimulus period. Alpha band power increased slightly in the early post-stimulus period, while decreasing slightly in the later post-stimulus period. As for oscillatory power, beta and low-gamma fractal power decreased, peaking between 400 and 600 ms. In contrast to the pattern in oscillatory power, high gamma fractal power decreased, with a similar time course to beta-band fractal power. These results and their implications will be discussed in greater detail in a forthcoming publication.

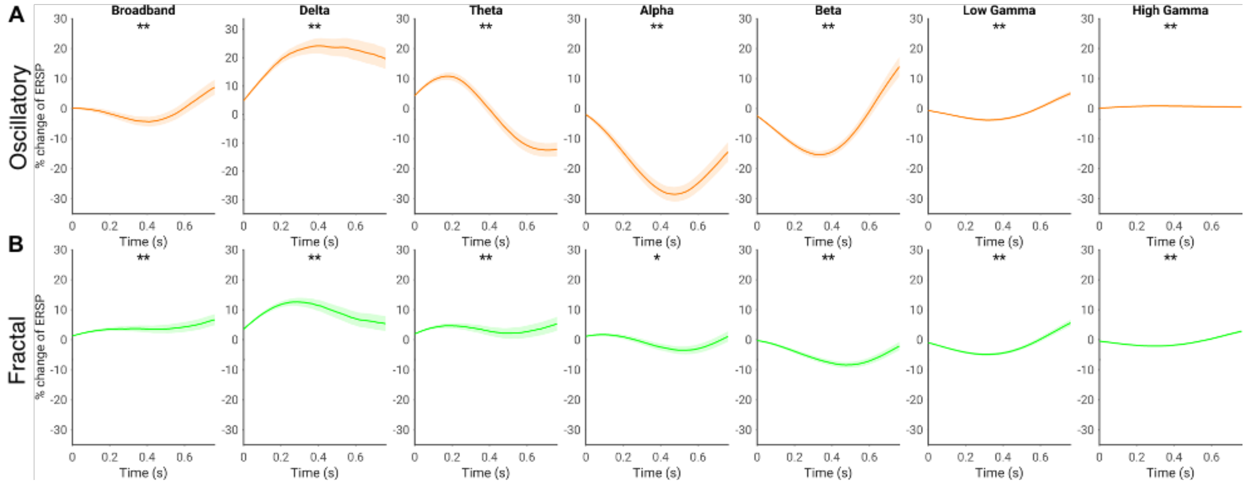

**Figure S1.** Spectral power response of oscillatory and fractal components. Asterisks indicate a significant difference from zero. Shading indicates standard error. * = $p < 0.05$, ** = $p < 0.01$.

*Validation of the pseudotrial and TTV methods in simulated negative-interaction and no-interaction scenarios – the effect of trial-varying signal-to-noise ratio*

It is possible that trial-by-trial variations in signal-to-noise ratio (SNR) could generate spurious evidence of a spontaneous-evoked interaction. This could occur as trials with high SNR will show a large evoked response, while trials with low SNR may show a smaller evoked response. We tested for this possibility using simulations, which are described in more detail in the methods.

Figure S2 shows the result of the additive simulation, in which no spontaneous-evoked interaction was present. Only one SNR regime showed a significant spontaneous-evoked interaction using the pseudotrial method. For 49 SNR conditions, the expected number of false positives is 2.45, so this is well within the expectation under chance conditions ($p = 0.7101$, binomial test). Furthermore, these interactions were small and confined to short periods of time, and do not qualitatively resemble at all the robust interactions observed in our data. Using the TTV method, eight SNR regimes showed significant decreases of TTV. This is significantly more than should be expected by chance alone ($p = 0.000648$, binomial test). It appears that the TTV method may be adversely affected by variations in signal-to-noise ratio, particularly in certain noise regimes; however, we note that the effects observed in this data are rather small in comparison to the TTV reductions observed in the empirical data. Several other conditions showed a significant increase in TTV in the post-stimulus period – we do not consider these results in the statistics, since, as discussed in the methods, an increase in TTV is to be expected in purely additive conditions and is not indicative of spontaneous-evoked interaction.

Figure S3 shows the result of the spontaneous-evoked interaction simulation, in which a negative interaction between spontaneous and evoked activity was introduced. Using the method of pseudotrials, 17 SNR conditions showed significant spontaneous-evoked interaction in the expected direction ($p = 9.82 \times 10^{-12}$, binomial test). A further two displayed limited evidence of spontaneous-evoked interaction in the opposite direction ($p = 0.2139$, binomial test with $n = 32$). These conditions were generally the high SNR and low SNR variability conditions, as should be expected. Using the method of TTV, 28 conditions showed significant TTV decreases ($p = 0$, binomial test). Again, these conditions usually had high SNR and low SNR variability, but TTV decreases were visible in lower SNR conditions than for the method of pseudotrials.

Taken together, these results suggest that the method of pseudotrials is a robust and accurate method for assessing spontaneous-evoked interaction, and is not subject to false positives due to time-varying noise. The method of TTV, by contrast, appears to have greater power to detect negative interaction than the pseudotrial method, but it occasionally exhibits false positives due to trial-by-trial variations in noise contamination.

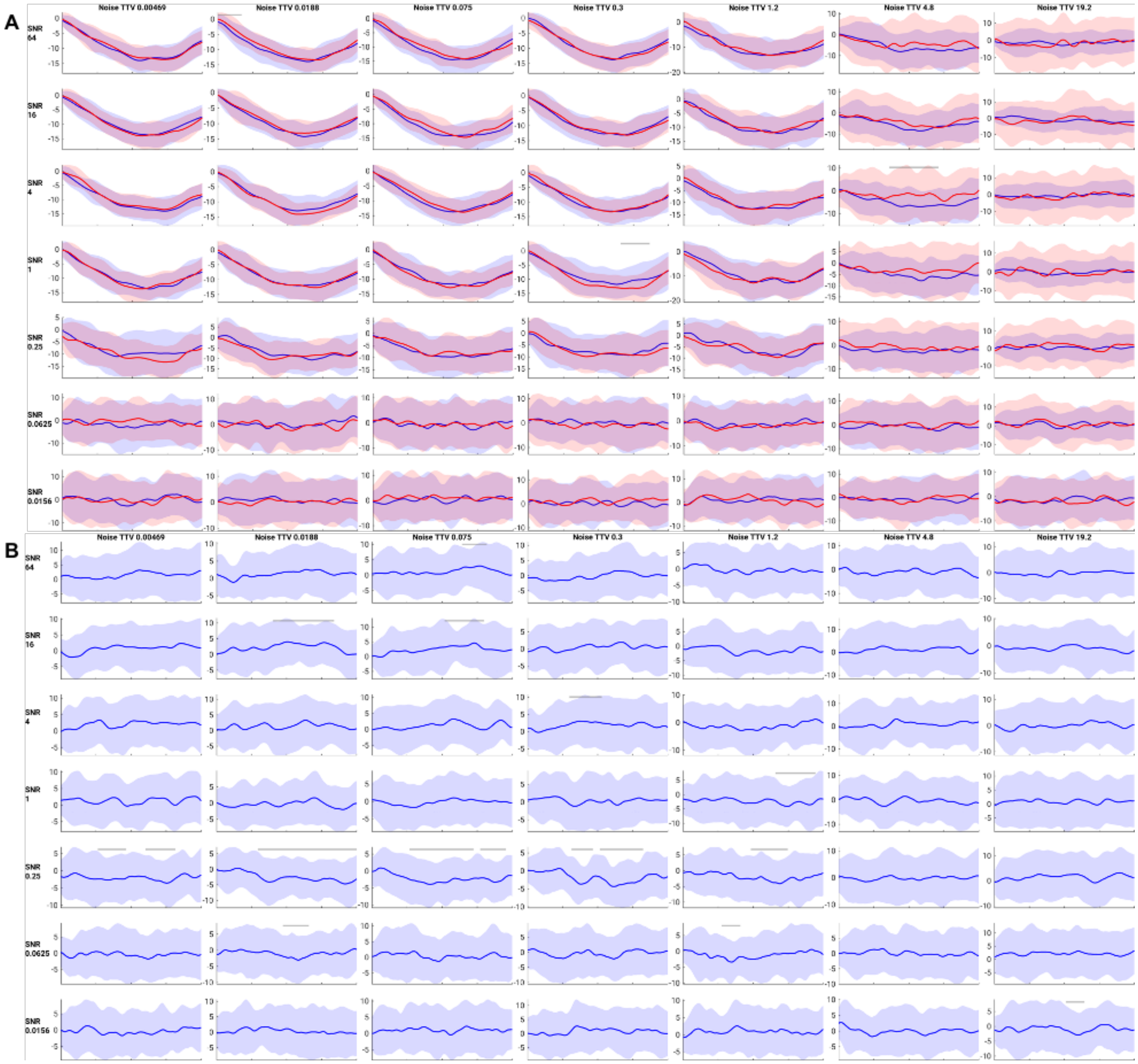


**Figure S2.** Results of the negative control simulation study, using both the method of pseudotrials and the method of trial-to-trial variability. A) Pseudotrial-based methodology. Rows indicate different average signal-to-noise ratios, and columns indicate different variabilities of noise across trials. Each plot shows the pseudotrial-corrected time course for the high prestimulus (red) and low prestimulus (blue) conditions. Shaded overbars indicate significance following a cluster-based permutation test (clustering across time only). B) TTV-based methodology. Each plot shows the TTV time course – shaded overbars indicate significance as in (A).

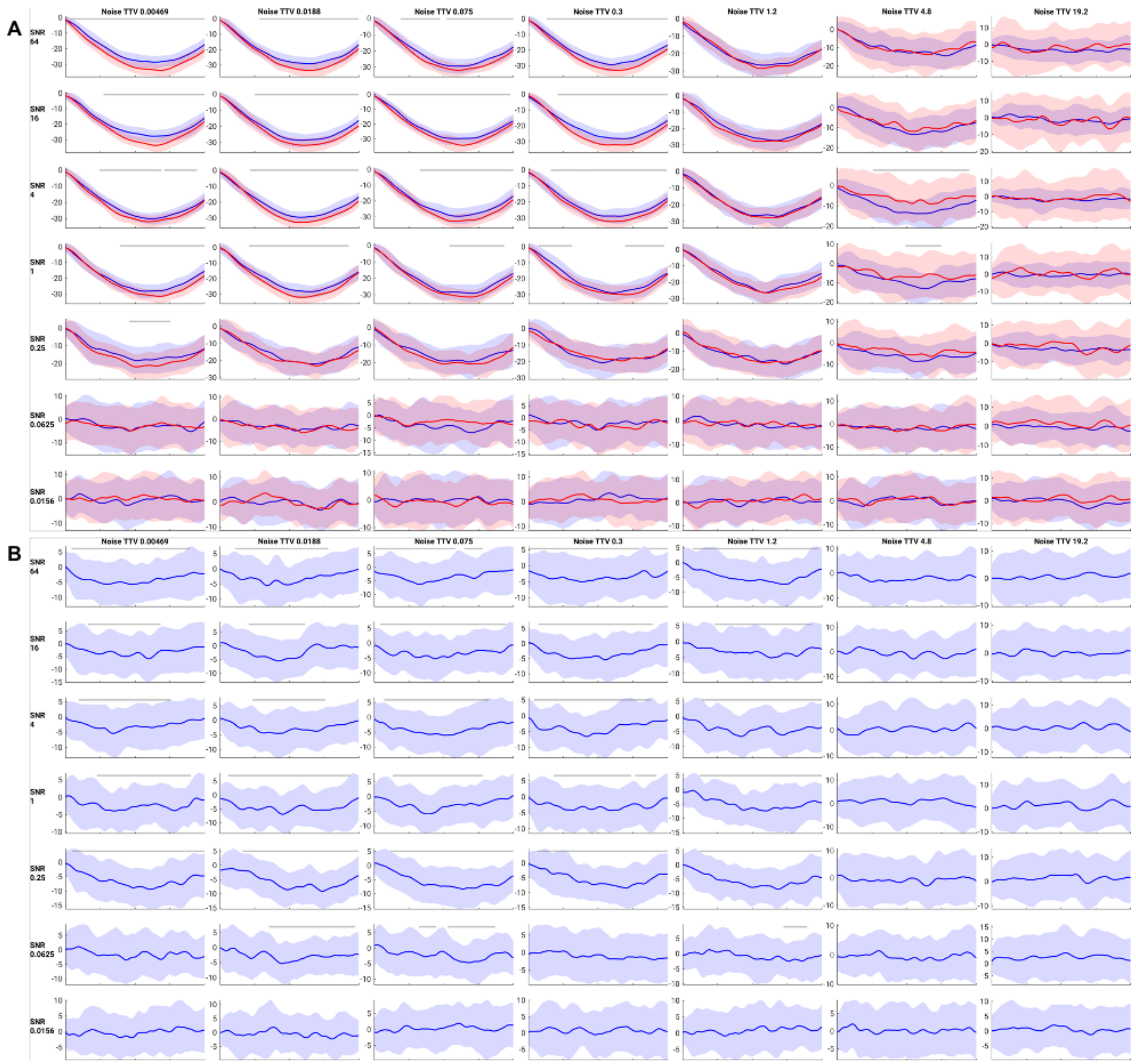

**Figure S3.** Results of the positive control simulation study, using both the method of pseudotrials and the method of trial-to-trial variability. A) Pseudotrial-based methodology. Rows indicate different average signal-to-noise ratios, and columns indicate different variabilities of noise across trials. Each plot shows the pseudotrial-corrected time course for the high prestimulus (red) and low prestimulus (blue) conditions. Shaded overbars indicate significance following a cluster-based permutation test (clustering across time only). B) TTV-based methodology. Each plot shows the TTV time course – shaded overbars indicate significance as in (A).

*Assessing the effect of prestimulus length – 50 and 200 millisecond prestimulus intervals*

We performed control analyses to ensure that the length of the prestimulus window chosen did not substantially affect the results of our analyses. Different frequency bands have different cycle durations, and as such it is possible that different lengths of the prestimulus period may affect the accuracy by which one can assess power within that window. We tried a 50 millisecond and a 200 millisecond prestimulus window, in addition to the 100 millisecond window in the main text, and replicated the analyses of figures 2 and 3 from the main text. The results were almost entirely consistent with the results of the main text.

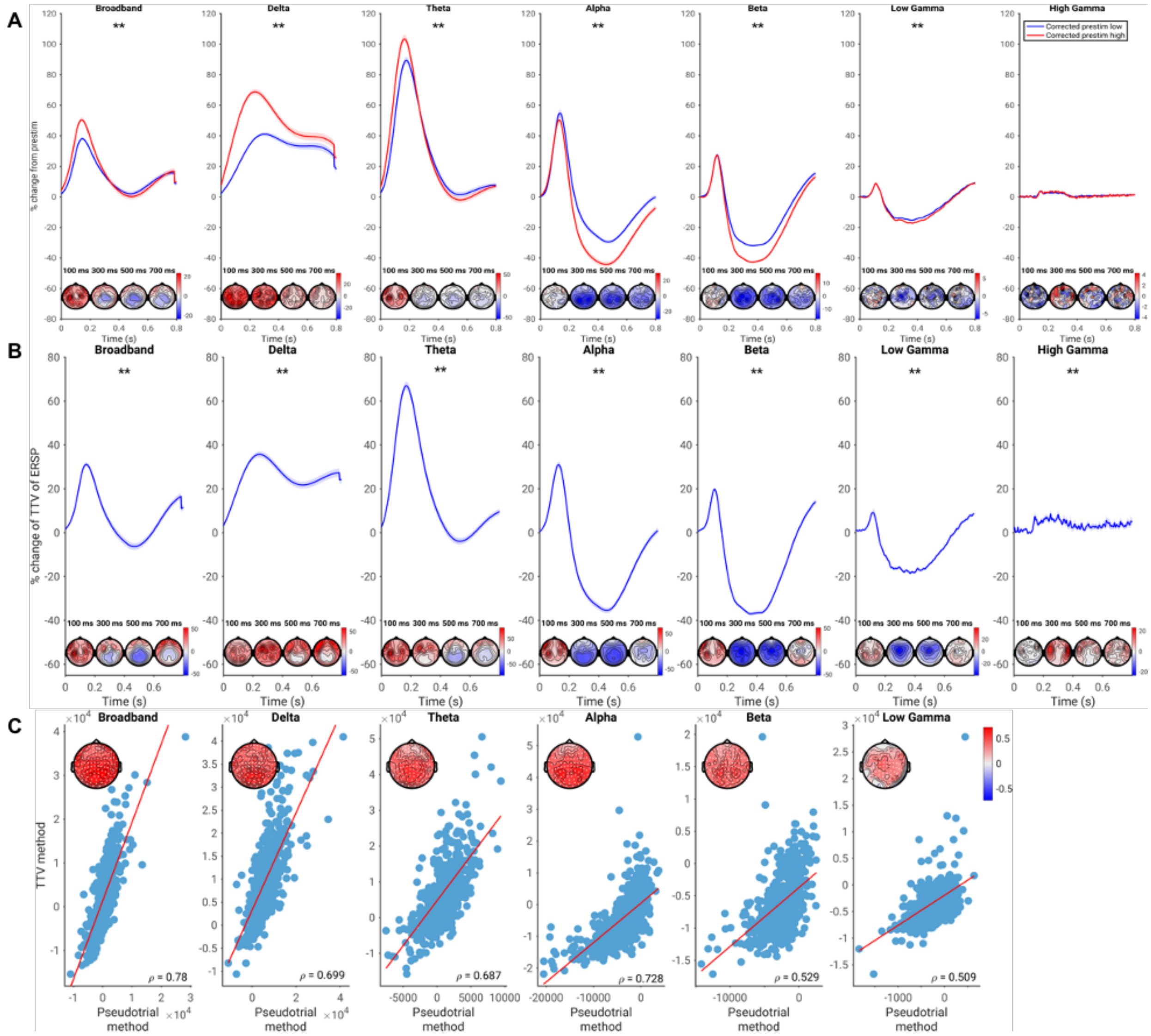


**Figure S4.** As figure 2 in the main text but using a 50 millisecond prestimulus window.

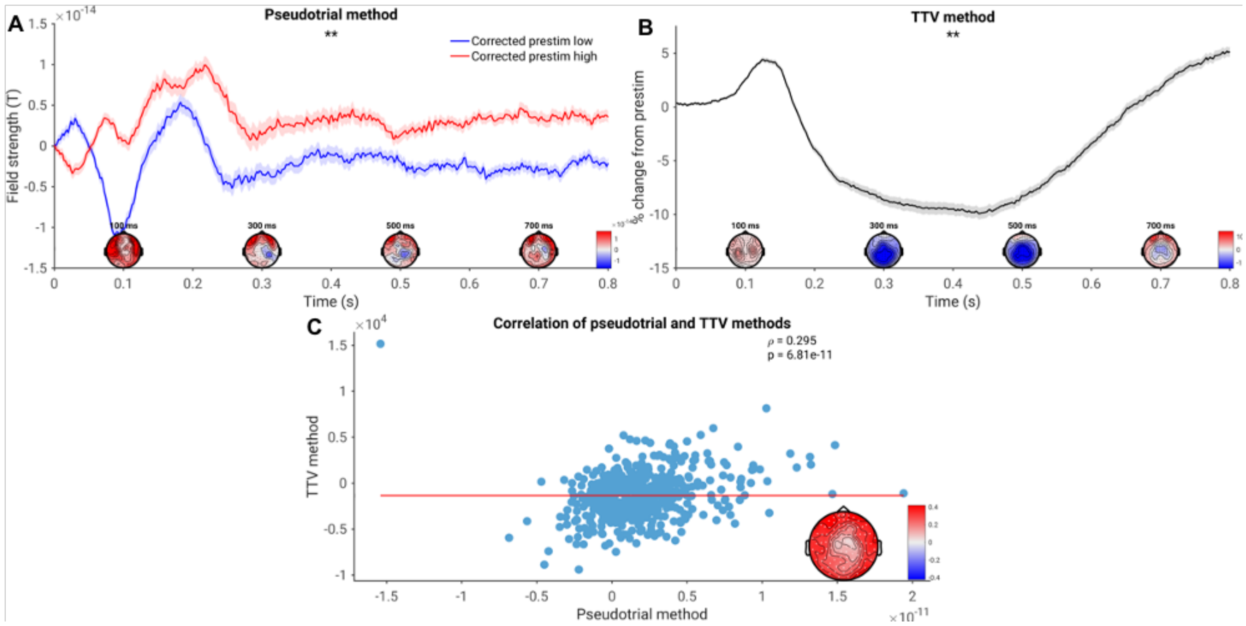


**Figure S5.** As figure 3 in the main text, but using a 50 millisecond prestimulus window.

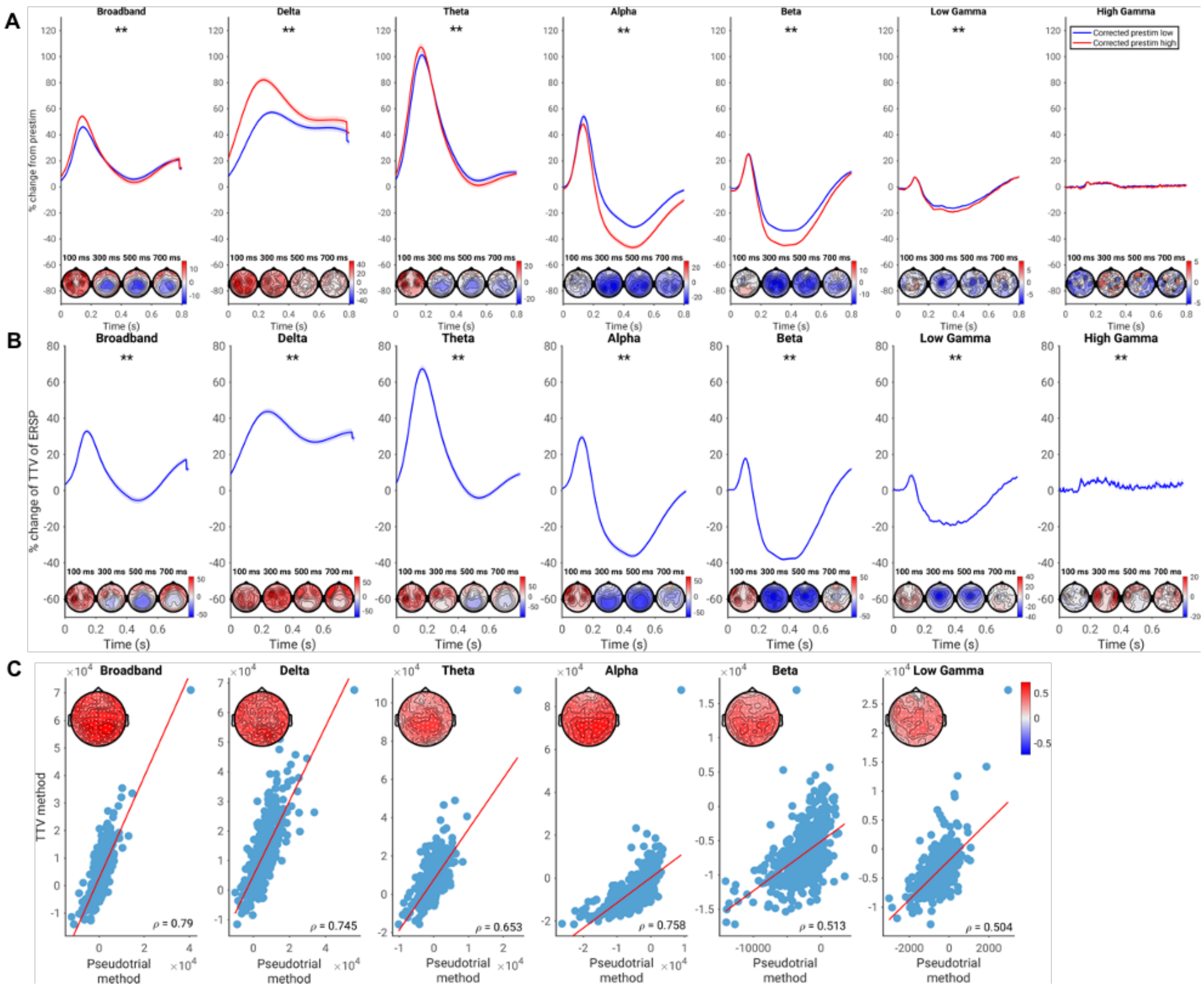


**Figure S6.** As figure 2 in the main text but using a 200 millisecond prestimulus window.

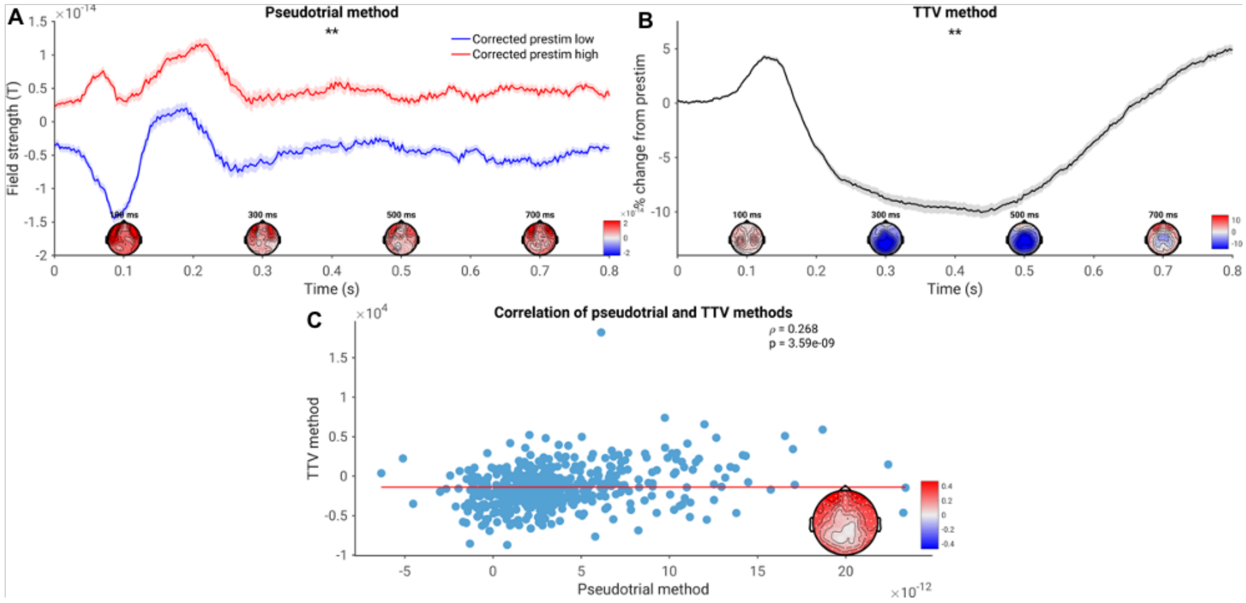


**Figure S7.** As figure 3 in the main text but using a 200 millisecond prestimulus window.

*Spontaneous-evoked interaction of oscillatory and fractal dynamics using the method of TTV*

In the main text, spontaneous-evoked interaction of oscillatory and fractal dynamics was assessed using the method of pseudotrials. Here we show that broadly analogous results hold using the method of TTV. For oscillatory power, we observed a TTV increase in delta ($p = 0.002$) and theta ($p = 0.002$), consistent with the positive interaction observed with the method of pseudotrials. The subsequent decrease in TTV in theta was not significant ($p = 0.745$). In alpha, beta, and low gamma, we observed a TTV decrease ($p = 0.002$ in each case), consistent with the negative interaction observed using the pseudotrial method. For fractal power, we observed a significant TTV increase in delta ($p = 0.002$) and theta ($p = 0.008$), consistent with the observations using the method of pseudotrials. However, we also observed a significant decrease in theta TTV in central sensors around 400 ms poststimulus ($p = 0.006$), indicating a negative interaction. Again contrary to the results using the method of pseudotrials, we observed a negative interaction in alpha ($p = 0.002$). Results were consistent with the method of pseudotrials for beta and low gamma, with both bands showing TTV decrease ($p = 0.002$). Finally, high gamma showed a negative interaction using the method of TTV ($p = 0.002$).

For most major effects, the TTV method and the pseudotrial method were in agreement for the oscillatory and fractal component data; however, there are a few notable differences. We submit that these differences may be due to overestimation of positive interaction using the method of pseudotrials (see figure S10 for details).

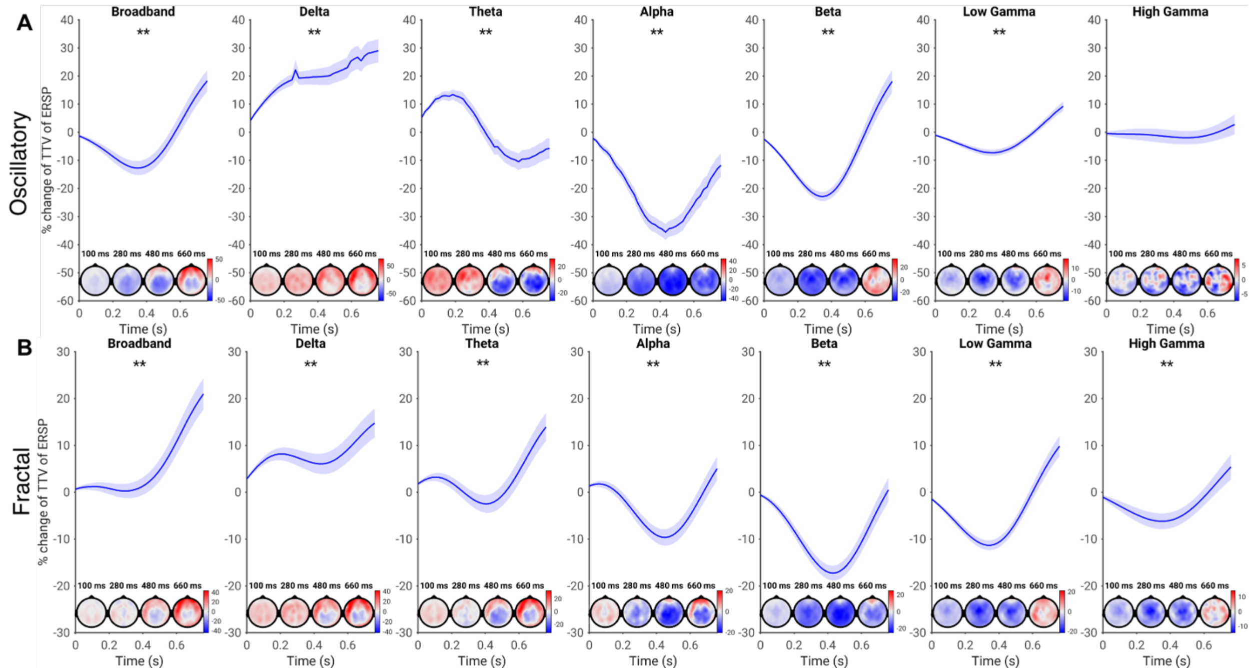


**Figure S8:** Spontaneous-evoked interaction of oscillatory (A) and fractal (B) power, assessed using the method of trial-to-trial variability. Shaded area indicates standard error. Effect topographies are shown at 100 ms, 300 ms, 500 ms, and 700 ms. * = $p < 0.05$, ** = $p < 0.01$.

*The effect of filtering on time-domain spontaneous-evoked interaction*

It is known that high-pass filtering at 1 Hz can distort event-related potential (ERP) waveforms, altering their latency and amplitude[3]. In the main text, we chose to use highpass filtering at 1 Hz because this filtering best facilitates ICA decomposition, reducing the level of artefact contamination in the main results[4]. We report the analyses of spontaneous-evoked interaction on the unfiltered data here, in order to assess whether highpass filter distortion had an effect on spontaneous-evoked interaction.

Preprocessing of this dataset was the same as described in the main text, but a copy of the data without the highpass filter was kept. The initial run of Autoreject, ICA training, and component classification were carried out on 1 Hz highpass-filtered data, but the weights from the ICA decomposition were applied back to the original, unfiltered data. Components classified as artefactual from this filtered data were then rejected in the unfiltered data. The remainder of preprocessing proceeded as described in the main text. n for this sample was slightly lower ($n = 440$ vs. 474) due to preprocessing errors.

Using the method of pseudotrials, we again noted a positive interaction in the time-domain signal ($p = 0.002$). However, in contrast to the main text results, this positive interaction was more limited in scope and magnitude, lasting only the first 200 milliseconds of the poststimulus period, rather than extending over the whole poststimulus period. Over central sensors, a negative interaction was observed between 300 and 400 milliseconds post-stimulus ($p = 0.002$; see topoplots). Results using the method of TTV were largely similar to the main text results: we observed a widespread decrease in TTV in the poststimulus period, as found in previous papers (Arazi et al., 2017a,b; $p = 0.002$) as well as a brief TTV increase at approximately 100 ms poststimulus (p = 0.01). The results of the two methods remained significantly correlated (p = 0.002), though this correlation remained small in magnitude.

These results suggest that 1 Hz highpass filtering may have exaggerated the magnitude of positive interaction in our data, and masked the presence of a topographically-limited negative interaction. This may have occurred if part of the ERP response was smeared backwards in time by the effect of filtering, as observed by[3]. This would have pushed part of the post-stimulus response into the pre-stimulus period, exaggerating the correlation between pre-stimulus and post-stimulus. We emphasize, however, that the finding of positive interaction remains significant even when this effect has been controlled for.

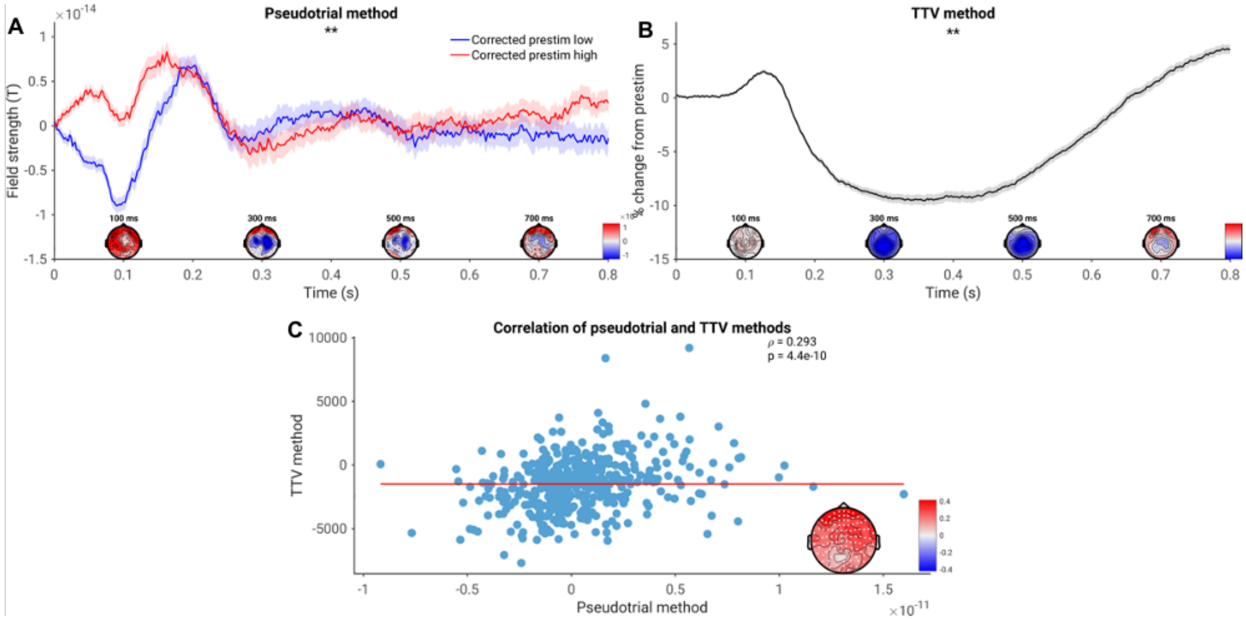


**Figure S9.** As figure 3 in the main text, but without the application of a 1 Hz highpass filter.

*The effect of temporal resolution on spontaneous-evoked interaction in low frequencies*

As discussed above in figure S9, any mechanism which results in part of the post-stimulus response being included in the pre-stimulus period may artificially inflate the magnitude of positive interaction observed. The estimation of low-frequency activity is inherently temporally imprecise, as the window size necessary to resolve low-frequency activity is large. As a result, it is possible that estimates of low-frequency activity in the prestimulus period are influenced by the poststimulus response.

We controlled for this with the following procedure. First, we visualized the time course of the power of the lowest-frequency band included in our data (the delta band, 2-4 Hz), and determined the point at which it begins to rise (figure A). We then chose our prestimulus period as the 100 milliseconds before this observable rise: conservatively, this was -400 ms to -300 ms. We then set the pseudotrial prestimulus period at the same offset from the pseudotrial. As a result, we were only able to analyze a 400 millisecond post-stimulus period: however, since low-frequency activity was of interest, this should be sufficient, as the low-frequency responses we observed occurred early in the post-stimulus period.

We observed that the magnitude of positive spontaneous-evoked interaction in the delta and theta bands was substantially reduced, from a maximum channel-average difference of 33% in delta to a maximum channel-average difference of 6%. However, the effect remained highly significant both in the delta ($p$ = 0.002) and theta bands ($p$ = 0.002). Negative interaction estimates were largely unaffected. These results suggest that the magnitude of positive spontaneous-evoked interaction in slow-frequency activity may have been overestimated due to the temporal imprecision inherent in resolving this activity. Once again, however, we note that positive interaction remained significant despite controlling for this effect.

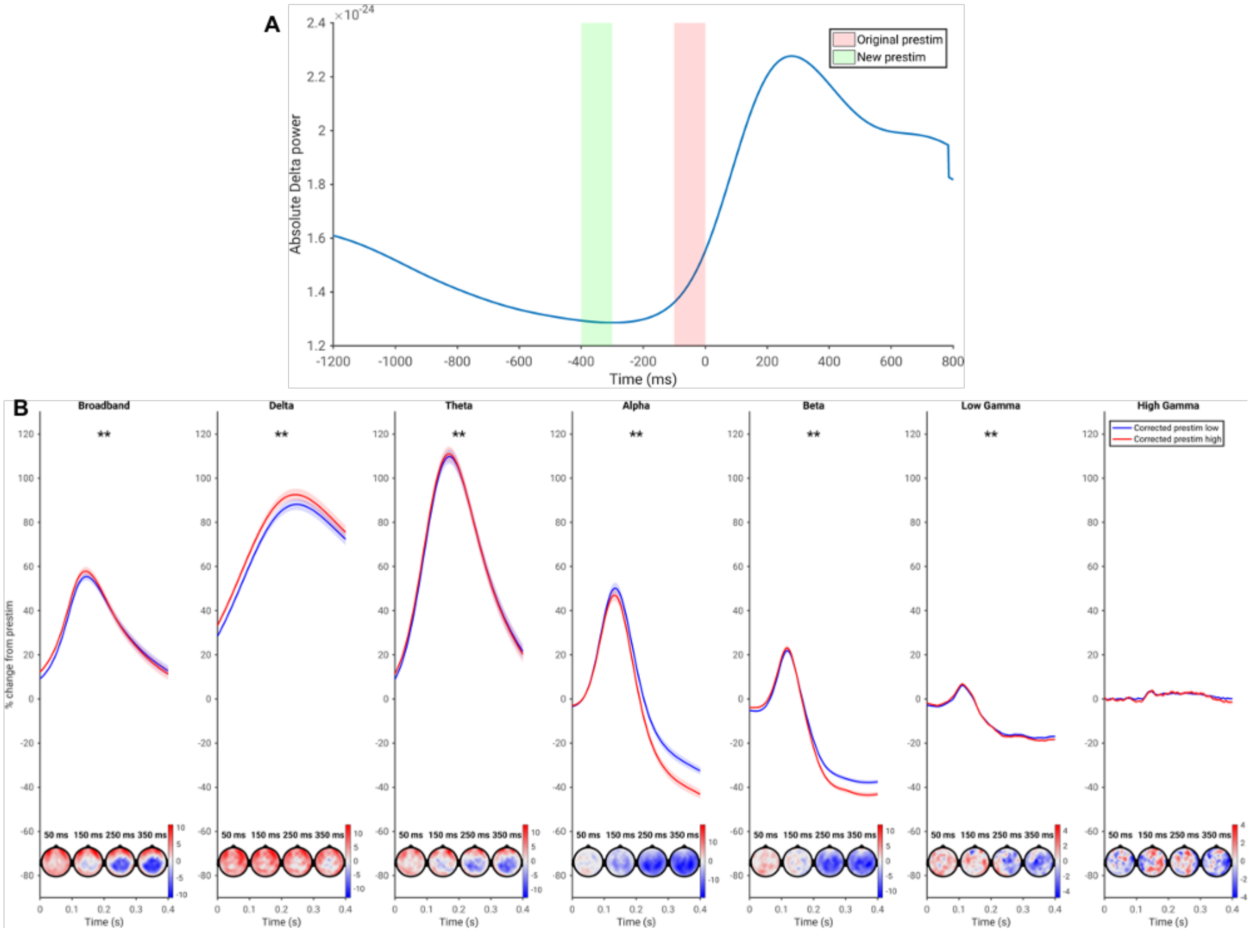


**Figure S10:** The effect of temporal resolution on low-frequency spontaneous-evoked interaction. A) The two prestimulus windows considered. The red highlight shows the original prestimulus window from the main text, while the green highlight shows the prestimulus window considered for panel B: note that this window falls before any increase in power is observed. B) Pseudotrial-corrected time courses and topoplots for analysis using the green prestimulus window from A. Asterisks, topoplots, and shading as in figure 2a) in the main text.

*Evaluating potential mechanisms for spontaneous-evoked interaction in the alpha band – the effect of phase-amplitude coupling*

Phase-amplitude coupling is a robust phenomenon observed in electrophysiological data in many settings, and between multiple pairs of frequency bands[5]. Phase-amplitude coupling has been associated with learning[6], working memory[7], and perception[8], and has been proposed as a mechanism for connecting large-scale network activity with local computation[5]. Additionally, phase synchronization of low-frequency activity is a phenomenon which has is frequently observed in the context of rhythmic stimulation[9,10], but also as a process of "phase resetting" following non-rhythmic stimuli[11,12].

In fMRI, a previous study showed that the magnitude of phase-amplitude coupling during resting-state predicted the degree of spontaneous-evoked interaction[13]. The combined effects of phase-coherence and phase-amplitude coupling are a potential mechanism for spontaneous-evoked interaction. If delta phase is coupled to alpha power, then post-stimulus delta phase synchronization could result in a change in alpha power. If phase-synchronization effects were driving this change in alpha power, then this change would be inherently non-additive. Phase synchronization implies some process of phase-resetting, which would result in different responses for different points in the prestimulus delta phase cycle. For example, assume that delta peaks synchronize in the poststimulus period, and the stimulus induces a "resetting" or "jumping" to the peak portion of the phase cycle. For trials where the prestimulus period occurred during the peak of a delta cycle, no change in power would be observed, as the same phase continues to the poststimulus period; on the other hand, for trials where the prestimulus period occurred during the trough of the delta cycle, a maximal change in power would be observed. In this way, the combination of inter-trial phase coherence and phase-amplitude coupling could produce a negative spontaneous-evoked interaction. Though not exhaustively or rigorously, we examined this possibility by assessing delta phase synchronization and its relation to the negative interaction observed in the alpha band.

Inter-trial coherence (ITC) was calculated as described in[14]. Phase-amplitude coupling (PAC) between the delta and alpha bands was assessed using the ERPAC[15]. We then correlated the magnitude of phase-amplitude coupling, the magnitude of inter-trial coherence, and their product with the spontaneous-evoked interaction observed in our data (assessed with the method of pseudotrials). We correlated the PAC and ITC measures with the pseudotrial-corrected difference between prestimulus high and low at each time point, correcting for multiple comparisons with a cluster-based permutation test across time and channels. We also assessed the relationship of PAC and ITC with spontaneous-evoked interaction in a static way: we created a summary index of PAC, ITC, and PAC*ITC by taking the area under the curve across all time points and sensors included in a significant cluster.

We observed significant phase-amplitude coupling at the group level ($p = 0.002$), as well as significant inter-trial coherence at the group level ($p = 0.002$). The former peaked at 300 ms and remained high, while the latter peaked at 250 ms and returned to baseline. No significant correlations were observed with PAC ($p = 0.5375$), ITC ($p = 0.1139$), or the interaction of the two ($p = 0.5315$) when considering the time-resolved correlations. Considering the summary index correlations, no significant correlations were observed for PAC ($p = 0.1499$) or ITC ($p = 0.3636$); a significant correlation was observed for PAC*ITC, but it was in the opposite direction as expected ($p = 0.0460$).

Though we did not investigate phase-amplitude coupling systematically as a mechanism for spontaneous-evoked interaction, these preliminary results indicate that at least for spontaneous-evoked interaction in alpha, delta-alpha phase-amplitude coupling is not a mechanism. Future research should investigate this hypothesis more thoroughly, as well as other potential mechanisms for spontaneous-evoked interaction.

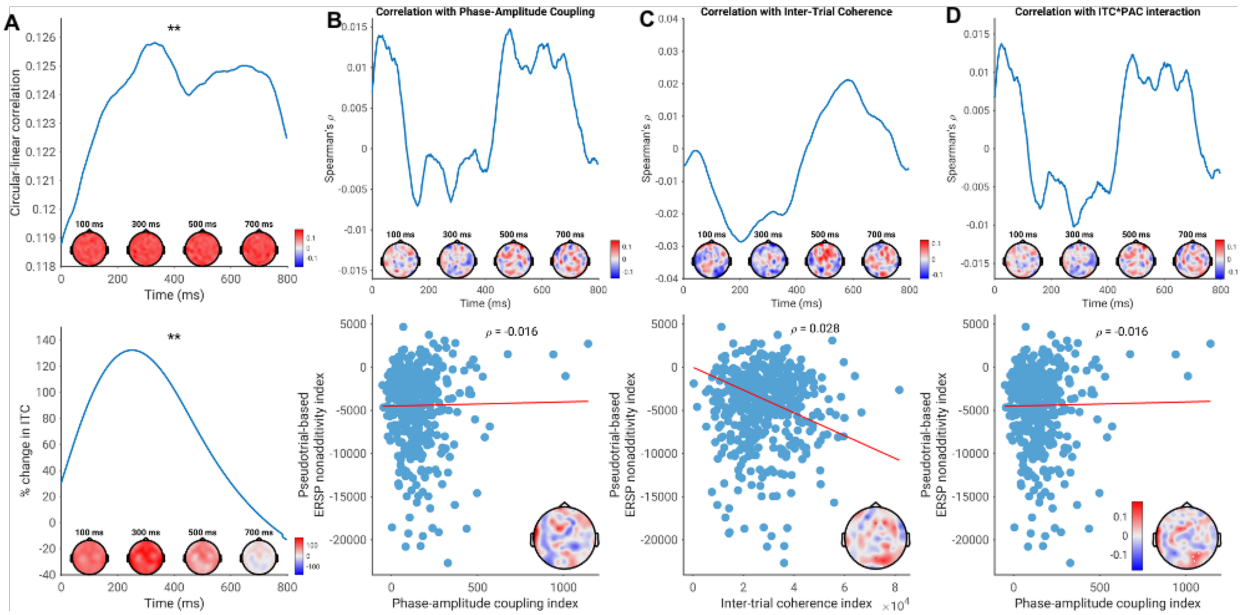


**Figure S11:** Effect of delta phase coherence and delta-alpha phase amplitude coupling on spontaneous-evoked interaction in the alpha band. A) (top) Phase-amplitude coupling between delta and alpha bands, averaged across subjects. Topoplots show the topography of the effect at the indicated latencies. Asterisks indicate significant difference from zero. * = p < 0.05, ** = p < 0.01. (bottom) Inter-trial coherence in the delta band, expressed as percent change from baseline. B) (top) Correlation at each time point between delta-alpha phase amplitude coupling and the magnitude of spontaneous-evoked interaction in alpha, assessed with the method of pseudotrials. Topoplots and asterisks as in A). (bottom) Correlation of summary indices of phase-amplitude coupling and spontaneous-evoked interaction. Scatter plots shows the correlation of the mean index across all sensors, while the topoplots show the topographical distribution of the correlation. White dots indicate sensors which are significant following a cluster-based permutation test. C) As B), but for inter-trial coherence instead of phase-amplitude coupling. D) As B), but for the interaction of PAC and ITC.
